# CellLoop: Identifying single-cell 3D genome chromatin loops

**DOI:** 10.1101/2025.04.08.647893

**Authors:** Yusen Ye, Yuxuan Hu, Shihua Zhang, Hebing Chen, Lin Gao

**Affiliations:** School of Computer Science and Technology, Xidian University, Xi’an 710126, Shaanxi, China; NCMIS, CEMS, RCSDS, Academy of Mathematics and Systems Science, Chinese Academy of Sciences, Beijing 100190, China; School of Mathematical Sciences, University of Chinese Academy of Sciences, Beijing 100049, China; Key Laboratory of Systems Health Science of Zhejiang Province, School of Life Science, Hangzhou Institute for Advanced Study, University of Chinese Academy of Sciences, Chinese Academy of Sciences, Hangzhou 310024, China; Institute of Health Service and Transfusion Medicine, Beijing 100850, China

**Keywords:** single-cell chromatin loops, loop frequency map, spatial domains, density-based peak sweeping

## Abstract

Single-cell 3D genome mapping offers insights into chromatin architecture, yet the inherent sparsity and high dimensionality of contact maps hinder reliable identification of chromatin loops at the single-cell level. Here, we present CellLoop, a single-cell chromatin loop detection algorithm based on a density-based center detection framework that integrates intra-cellular and neighboring inter-cellular contacts via a re-voting strategy. Applied to Dip-C data from the mouse brain, CellLoop demonstrates superior accuracy based on the genome spatial distance and compartment signals at the single-cell level. It reveals cell-specific chromatin loops linked to transcriptional regulation and improved cell state classification. In HiRES embryogenesis data, CellLoop refines cell subtype definitions by minimizing cell cycle effects. Integrating GAGE-seq with MERFISH in mouse cortex, CellLoop redefines spatial domain functions through chromatin loop dynamics. Together, CellLoop enables robust, scalable, and biologically meaningful single-cell chromatin loop detection.

## Introduction

Single-cell 3D genome technologies have rapidly advanced, enabling the characterization of chromatin architecture underlying cellular processes at single-cell resolution^1-8^. Analytical frameworks have uncovered key structural features, including specific chromatin interactions^3, 9^, topological associating domains (TADs)^10, 11^, and compartments^12, 13^, enabling the establishment of a relation between chromatin structures and transcriptional control^14^. However, due to the inherent sparsity and ultra-high dimensionality of single-cell contact maps, robust methods for identifying chromatin loops at the single-cell level remain lacking.

Recent efforts have begun to address this gap. For example, SnapHiC identifies chromatin loops by aggregating data across hundreds of cells^15^, and SimpleDiff quantifies interaction strength based on 3D spatial distances reconstructed from individual-cell genomes^3^. Yet SnapHiC relies on small cell populations and cannot capture cell-specific features, while SimpleDiff lacks the resolution to distinguish loops from general chromatin interactions. To our knowledge, no existing method allows for de novo identification of chromatin loops in individual cells.

Here, we introduce CellLoop, an algorithm for single-cell loop detection based on a density-based center detection framework^16^. CellLoop integrates two complementary signals: (i) intra-cellular topology, which captures the spatial proximity of genomic loci within a single cell, and (ii) inter-cellular background strength, which reflects interaction probabilities across neighboring cells in a defined biological context. The algorithm evaluates each contact in parallel using two simple metrics, enabling efficient loop detection across thousands of single cells.

We benchmarked CellLoop using two orthogonal measures: genome spatial distance (GSD) and compartment property consistency (CPC) between loop anchors, demonstrating superior accuracy in single-cell level. Furthermore, we show that CellLoop can generate a loop frequency map (LFmap) to represent chromatin loop prevalence across cells, demonstrating improved accuracy and biological interpretability over existing approaches. By applying CellLoop to diverse single-cell 3D genomic datasets, we demonstrate its ability to uncover chromatin loop-based cell signatures and to refine cell state definitions and spatial domain functions. Together, these findings highlight the utility of chromatin loops as interpretable and biologically meaningful features in single-cell genome architecture analysis.

## Results

### 1. Overview of CellLoop algorithm

CellLoop efficiently identifies chromatin loops in individual cells using a density-based center detection algorithm^16^, which integrates intra-cellular and neighboring inter-cellular contact maps through a re-voting strategy (Fig. 1a). The method is applicable for generating LFmap and annotating cell states from the perspective of chromatin loops. It can be applied to various types of single-cell 3D genome datasets, enabling refined cell state identification and linking chromatin loops to specific cellular functions (Fig. 1b).

**Fig. 1:**
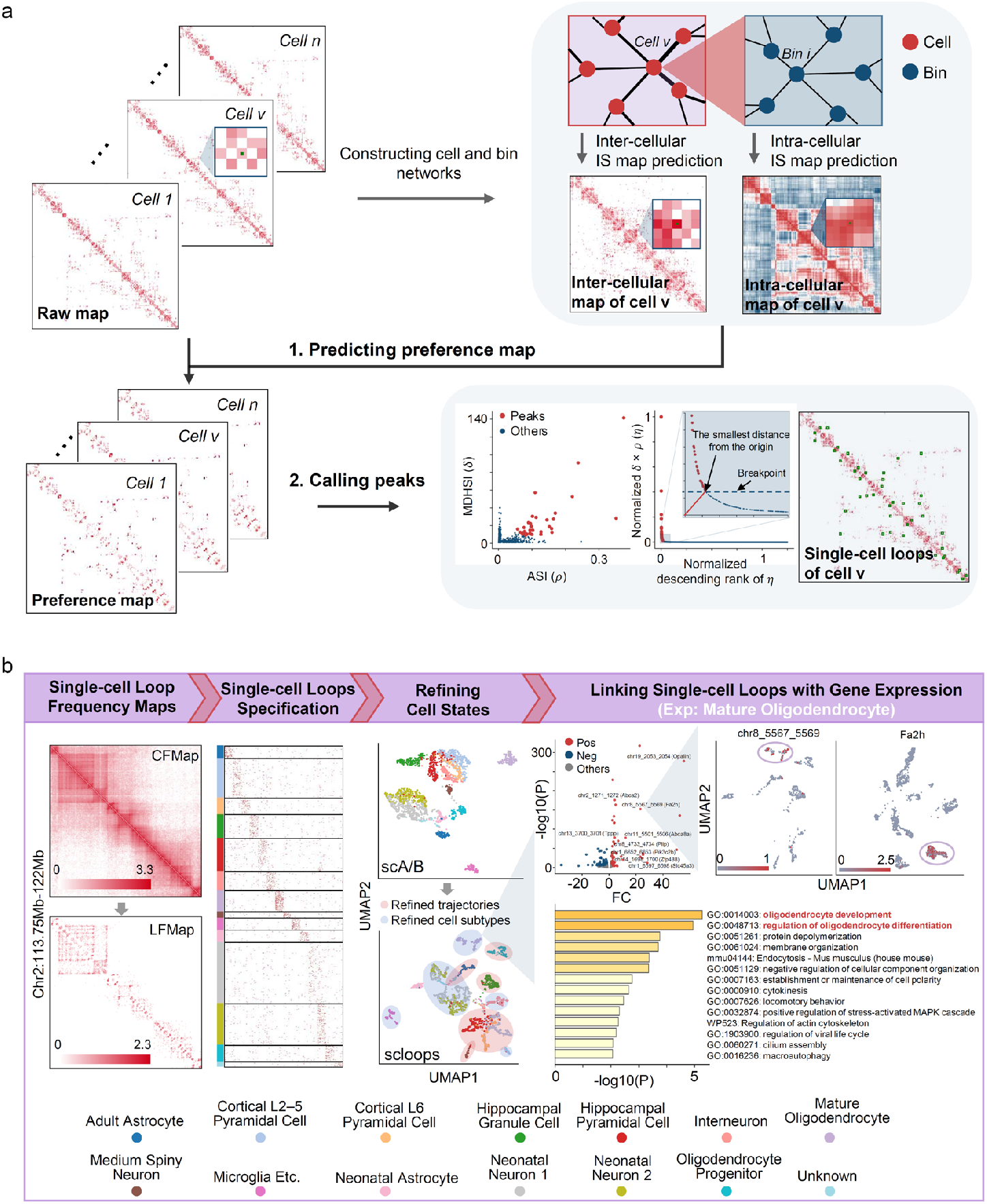
Overview of the CellLoop Algorithm. **a**, Schematic diagram of the CellLoop algorithm. CellLoop comprises two main components: Preference prediction module and Peaks Calling module. **Module 1:** CellLoop initiates the process by constructing cell and bin networks using raw contact maps and initial cell embeddings. It then generates interaction strength (IS) maps at both inter-cellular and intra-cellular levels. Based on these maps, CellLoop predicts a preference map that governs chromatin interaction formation within a specific single cell. **Module 2:** For each candidate chromatin interaction, CellLoop defines two metrics: the aggregated strength of chromatin interactions (ASI, *ρ*) and the minimum distance of higher-strength chromatin interactions (MDHSI, *δ*). CellLoop ranks the overall chromatin interaction score (*η* = *ρ*×*δ*) in descending order to infer breakpoints and differentiate chromatin loops from candidate interactions. **b**, The significance of CellLoop in the Dip-C dataset is highlighted in the following aspects. The first column, the Chromatin Loop Frequency Map (LFmap), shows greater interpretive power compared to the Chromatin Contact Frequency Map (CFmap). The second column demonstrates how CellLoop detects cell-type-specific chromatin loop features. The third column illustrates that single-cell chromatin loop features (scloops) more effectively define cell states compared to other single-cell 3D features. The fourth column shows how CellLoop links cell-type-specific chromatin loops with gene expression in mature oligodendrocytes.

CellLoop consists of two primary components (Fig. 1a): 1. **Preference prediction module:** The input to CellLoop includes raw contact maps and an initial cell embedding (Methods), These are used to separately construct intra-cellular and inter-cellular networks. In the intra-cellular network, bins represent nodes, and edge weights are determined by contact frequencies between bins, reflecting the strength of intrachromosomal topological associations within individual cells. The inter-cellular network, where cells are nodes and edge weights are based on the correlation coefficients of cell embedding vectors, captures the strength of chromatin interactions within a specific biological context. The combined data from varying support strengths and raw contact maps together inform the preference of chromatin interaction formation in a given cell. 2. **Peaks calling module:** Once the preference map of chromatin interactions is obtained for a single cell, CellLoop formulates the problem as peak calling from a two-dimensional matrix and applies the center detection algorithm to identify chromatin loops in that cell.

### 2. Performance evaluation of CellLoop

We applied CellLoop to Dip-C data generated from the developing mouse cortex and hippocampus at a 20Kb resolution^2^. CellLoop identified over 260,000 chromatin loops occurring in at least five cells, with a median of 1,769 chromatin loops per cell (Fig. 2a). Of these loops, more than 98% were detected in fewer than 64 cells, while 1,387 loops were detected in over 128 cells (Fig. 2b). CellLoop detected a median of 816.5 chromatin loops per cell at genomic distances <80Kb, and 357 loops per cell at distances ranging from 320Kb to 640Kb. In contrast, only a median of 5 chromatin loops were detected at distances >2.56Mb (Fig. 2c). Additionally, we found that 42.3% of the chromatin loops identified by CellLoop contained at least one gene within their two anchors (Fig. 2d).

**Fig. 2:**
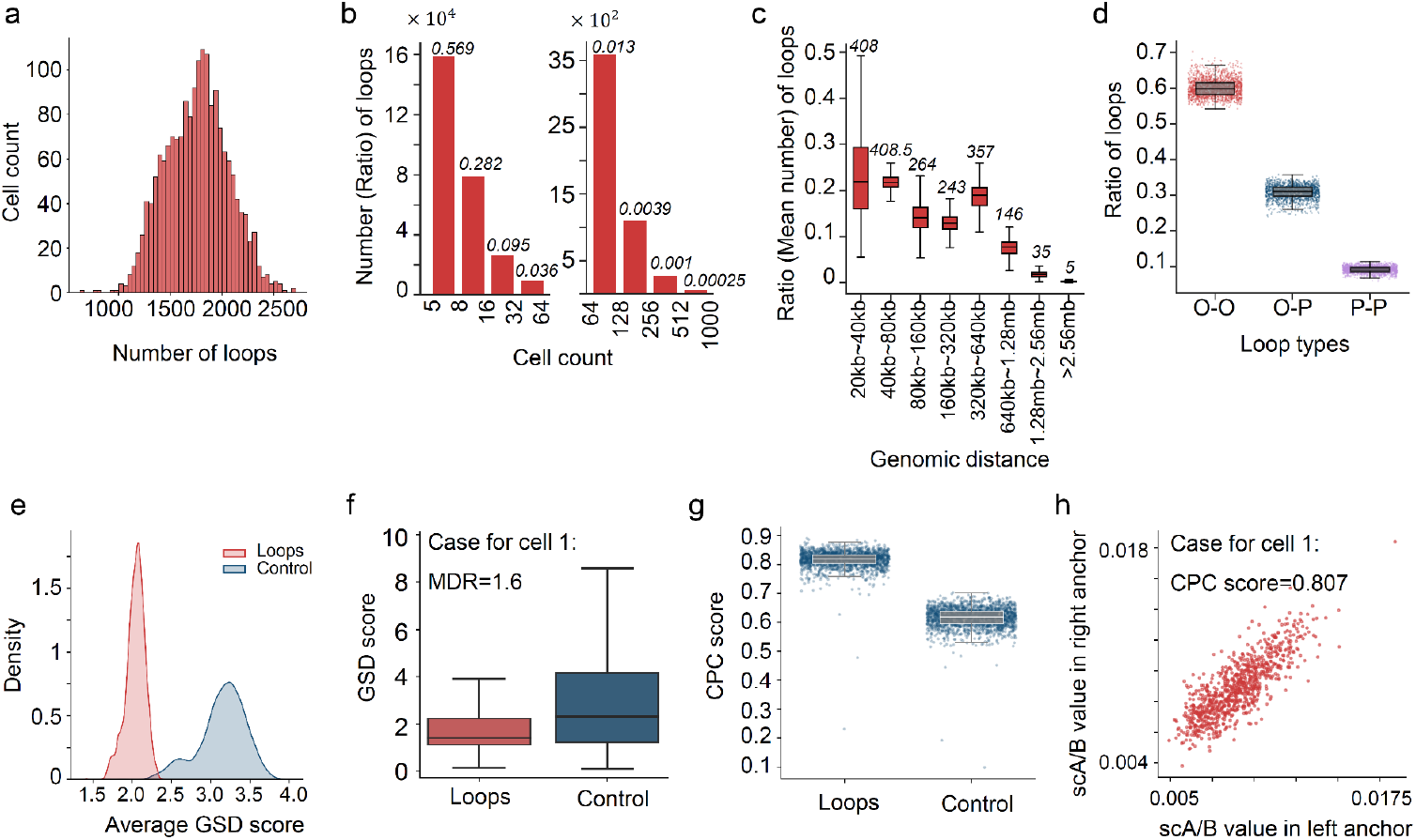
Performance evaluation of CellLoop algorithm in the Dip-C dataset. **a**, Distribution of chromatin loop numbers across different single cells. **b**, The number (and ratio) of chromatin loops detected in varying numbers of cells. For example, nearly 160,000 chromatin loops (accounting for 0.569) were detected simultaneously in 5-8 cells. **c**, Distribution of single-cell chromatin loops across different genomic distances. **d**, Proportion of different types of chromatin loops in single cells. **e**, Density distribution of the average genome spatial distance (GSD) scores for chromatin loops and their control interactions in single cells. **f**, GSD scores of chromatin loops and their control interactions in cell 1. **g**, Distribution of compartment property consistency (CPC) scores for chromatin loops and their control interactions in single cells. **h**, Distribution of scA/B values for the two anchors of chromatin loops and CPC scores for cell 1.

To assess CellLoops performance, we evaluated two metrics: genomic spatial distance (GSD) and compartment property consistency (CPC) between the anchors of chromatin loops in individual cells (Methods). Authentic chromatin loops are expected to exhibit lower GSD and higher CPC scores compared to control interactions. Consistent with this expectation, CellLoop-identified chromatin loops demonstrated significantly reduced GSD scores relative to controls (Fig. 2e). In cell 1, for instance, the mean GSD score of control interactions was 1.6 times that of the identified loops (Fig. 2f). Furthermore, the identified loops displayed markedly higher CPC scores than controls (Fig. 2g), with cell 1 achieving a CPC score of 0.807 (Fig. 2h). These findings indicate that CellLoop effectively characterizes single-cell chromatin loop features through both GSD and CPC metrics.

To demonstrate CellLoops efficiency, we implemented parallel processing utilizing 10 CPU cores. Notably, CellLoop identified single-cell chromatin loops for a single chromosome in an average of 0.99 seconds at 100Kb resolution and approximately 13 seconds at 10Kb resolution, enabling the analysis of tens to hundreds of thousands of single-cell datasets within a reasonable timeframe (Supplementary Fig. 1 and Supplementary Notes).

### 3. Performance evaluation of CellLoop by LFmap

We further merged chromatin loops of single cells detected by CellLoop to generate LFmap for the developing mouse cortex and hippocampus. LFmap offers several advantages over the traditional contact frequency map (CFmap): 1. **Reduced genomic distance bias:** LFmap effectively diminishes biases associated with genomic distance. The correlation coefficient between contact frequency and genomic distance in CFmap ranges from 0.77 to 0.84 across various chromosomes, whereas in LFmap, this value decreases to between 0.28 and 0.35 (Fig. 3a). 2. **Enhanced detection sensitivity:** Benchmarking against the two most popular methods, including a commonly used loop detection method for bulk 3D genome map, HiCCUPs^17^, and a loop detection method for single-cell 3D maps, SnapHiC^15^. LFmap demonstrates superior sensitivity. For chromosome 2, the aggregated peak score (APS) from LFmap is over four times higher than that from CFmap for chromatin loops identified by both HiCCUPs and SnapHiC (Fig. 3c,d). LFmap identifies over 52,000 chromatin loops present in at least three single cells, covering 92.7% of genomic regions, compared to 1,192 loops (11.4% of regions) by SnapHiC and 278 loops (5.3% of regions) by HiCCUPs (Extended Data Fig. 1a,b). 3. **Comprehensive integration:** LFmap consolidates chromatin loops detected by various computational methods^18^. Only 54 chromatin loops are commonly identified by both HiCCUPs and SnapHiC. LFmap encompasses 50 of these with high enrichment (*P* value< 10^−5^ in Fig. 3e,f and Supplementary Notes). On average, LFmap covers 88.4% of loops detected by both methods, 69.5% of those identified solely by SnapHiC, and 75% by HiCCUPs alone across all chromosomes (P value approaching 0 from the hypergeometric test; Extended Data Fig. 1c and Supplementary Notes). Notably, loops overlapping with LFmap exhibit significantly higher interaction frequencies than non-overlapping ones (Extended Data Fig. 1d). 4. **Extended genomic distance coverage:** LFmap identifies chromatin loops across a broader range of genomic distances, particularly those less than 100Kb (Fig. 3g and Extended Data Fig. 1e). 5. **Enrichment in regulatory elements:** After excluding loops recognized by SnapHiC or HiCCUPs, the remaining LFmap loops are enriched for transcription and regulatory factors such as CTCF, H3K4me1, H3K4me3, H3K27ac, and Pol II (Fig. 3h and Methods). Loops enriched with CTCF predominantly span 400Kb to 1.6Mb, aligning with known CTCF/cohesion-linked loops^19^, while those associated with enhancer or promoter markers are mainly within 100-200Kb, consistent with transcription-linked loops^20^.

**Fig. 3:**
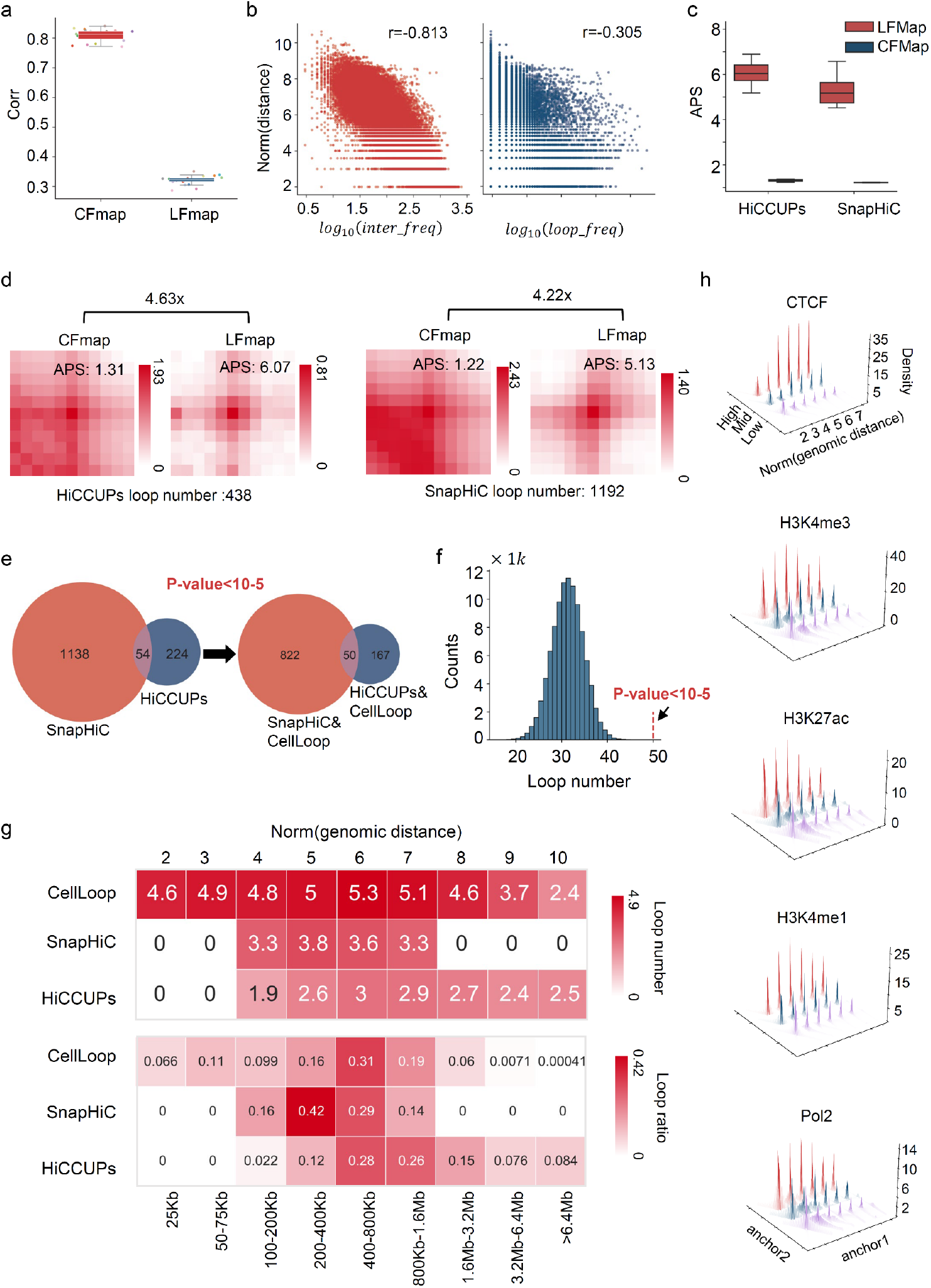
Performance evaluation of CellLoop by LFmap in the Dip-C dataset. **a**, Comparison of correlation coefficients between interaction frequency and genomic distance on the CFmap and LFmap across different chromosomes. **b**, Left: Scatter plot of interaction frequency versus genomic distance for interactions on chromosome 2 in the CFmap. Right: Scatter plot of interaction frequency versus genomic distance for interactions on chromosome 2 in the LFmap. **c**, Distribution of the aggregate peak score (APS) for chromatin loops identified by HiCCUPs and SnapHiC on the CFmap and LFmap, respectively, across all chromosomes. **d**, Comparison of APS for chromatin loops identified by HiCCUPs and SnapHiC on the CFmap and LFmap for chromosome 2. **e**, Left: Overlap of chromatin loops identified by SnapHi-C and HiCCUPs. Right: Overlap of chromatin loops detected by SnapHi-C & CellLoop and HiCCUPs&CellLoop for chromosome 2 (HiCCUPs&CellLoop represents chromatin loops identified by both methods simultaneously). **f**, Expected distribution of chromatin loop numbers in the LFmap. These loops are detected simultaneously by SnapHi-C and HiCCUPs. Statistical significance (*P* value) was determined by comparing the observed overlap count to the overlap counts generated from 100,000 random repetitions. **g**, Distribution of chromatin loop numbers (and their ratios) at different genomic distances for three methods. **h**, Enrichment distribution of ChIP-seq peaks around chromatin loops present in the LFmap but not identified by either SnapHiC or HiCCUPs.

In summary, LFmap offers enhanced biological interpretative power from the perspective of chromatin loops compared to previous methodologies.

### 4. Linking cell type-specific chromatin loops with gene functions

To elucidate the biological significance of single-cell chromatin loops, we utilized chromatin loops identified by CellLoop as features for UMAP visualization. This analysis demonstrated that, compared to scA/B features, single-cell chromatin loop features more distinctly classified cell types and identified finer cell subtypes (Fig. 4a).

**Fig. 4:**
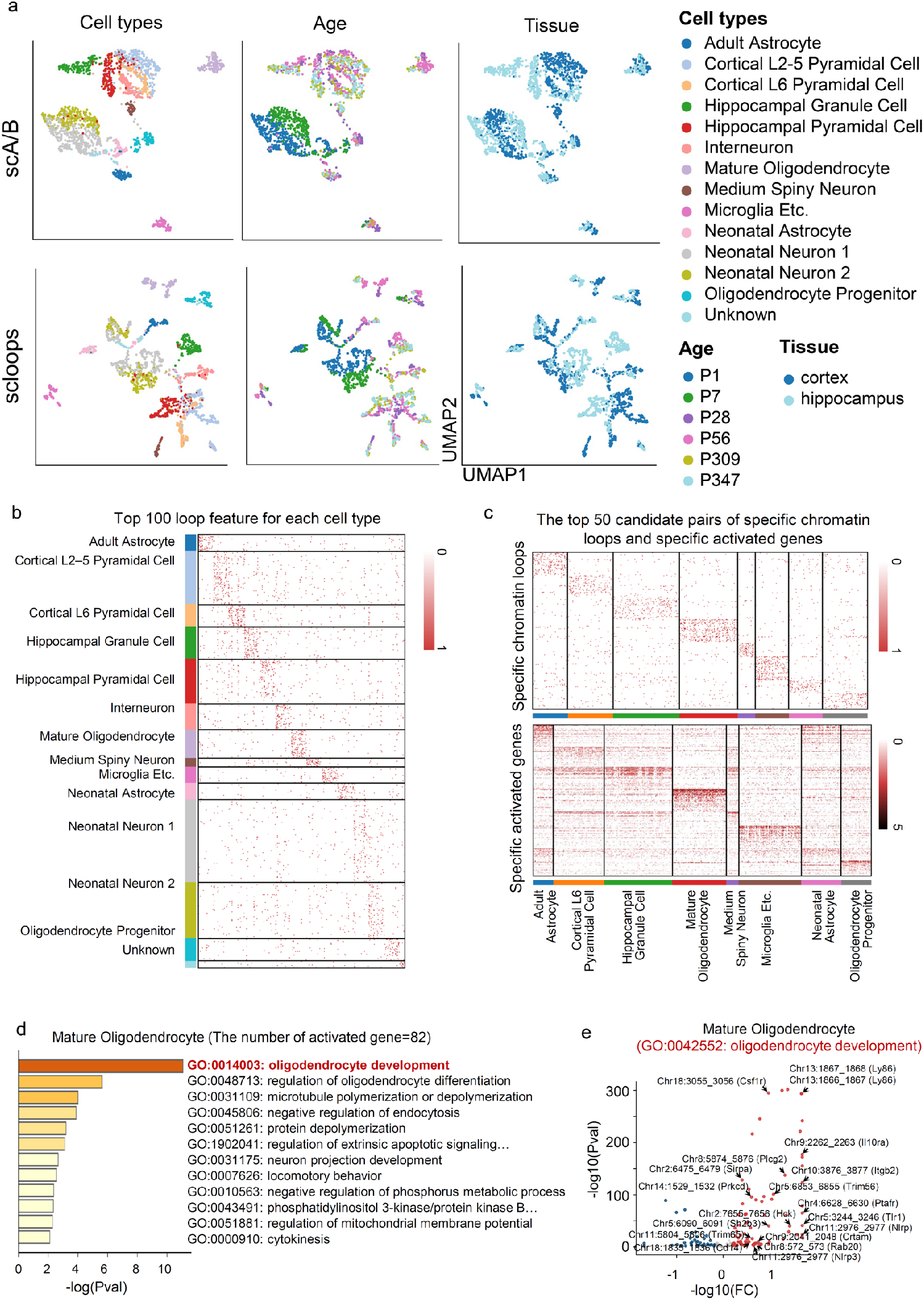
Application of CellLoop to the Dip-C dataset for linking cell type-specific chromatin loops with gene functions. **a**, UMAP visualization of cells using scA/B values and single-cell chromatin loop (scloops) features at 100KB resolution. Cells are labeled by cell type, age, and tissue from left to right. **b**, The top 100 most significant chromatin loops for each cell type, excluding the “Unknown” category. **c**, The top 50 Indi-Spe-highgene pairs for each cell type. **d**, Gene function enrichment analysis for gene sets from the Indi-Spe-highgene list of the Mature Oligodendrocyte cell type. **e**, Volcano plot of Indi-Spe-highgene pairs for the GO:0042552 term in the Mature Oligodendrocyte cell type.

We developed a computational pipeline to categorize detected chromatin loops into four subtypes (Extended Data Fig. 2 and Methods): 16,130 individual cell type-specific chromatin loops (Indi-Spe), 979 multiple cell type-specific chromatin loops (Multi-Spe), 1,402 common chromatin loops (Com), and 25,655 other chromatin loops (Others) (Extended Data Fig. 3a). Visual examples of Indi-Spe, Multi-Spe, and Com chromatin loops are provided in Extended Data Fig. 3b-d. We observed that the genomic distances between anchors of specific chromatin loops (median of 60 kb for both Indi-Spe and Multi-Spe) were shorter than those of Com loops (median of 100 kb) (Extended Data Fig. 3e). Additionally, Indi-Spe chromatin loops appeared in fewer cells than Com, Multi-Spe, and Other chromatin loops (Extended Data Fig. 3a). Across 13 cell types, we identified 465-2,135 Indi-Spe chromatin loops, with 30 Indi-Spe chromatin loops associated with unclassified types in the Dip-C data^2^ (Fig. 4b and Extended Data Fig. 3f). Notably, chromatin loops identified in non-neuronal cell types exhibited greater significance than those in neuron-specific cell types (Extended Data Fig. 3f).

Our pipeline further linked chromatin loops to specific gene expressions (Extended Data Fig. 2). By analyzing Indi-Spe chromatin loops alongside gene expression data from eight corresponding cell types in the Dip-C and MALBAC-DT^21^ datasets (Extended Data Fig. 4a,b), we identified 200-953 Indi-Spe chromatin loops containing at least one gene in their anchor regions (termed Indi-Spe-gene chromatin loops) and 103-449 Indi-Spe-gene chromatin loops with specific gene expression in their anchor regions for each cell type (Extended Data Fig. 3g). These chromatin loops were categorized into Indi-Spe-highgene and Indi-Spe-lowgene groups. For each cell type, the top 50 pairs within the Indi-Spe-highgene category are presented (see Extended Data Fig. 2 and Methods). The results revealed that both these chromatin loops and their associated genes exhibit distinct cell type specificity (Fig. 4c). For example, the chromatin loop at Chr8:5567_5569 is associated with the Fa2h gene in mature oligodendrocytes (Extended Data Fig. 4c,d), and the chromatin loop at Chr2:6475_6479 is linked to the Sirpa gene in microglia (Extended Data Fig. 4e,f).

Gene function enrichment analysis was performed for gene sets from Indi-Spe-highgene and Indi-Spe-lowgene categories of each cell type. In the Indi-Spe-highgene category, specific gene sets were enriched for Gene Ontology terms associated with specialized cell types, such as “innate immune response” in microglia and “oligodendrocyte development” in mature oligodendrocytes. In the Indi-Spe-lowgene category, relevant terms included “response to metal ion” in microglia and “neural retina development” in mature oligodendrocytes (Fig. 4d,e, Extended Data Fig. 4g,h, and Supplementary Figs. 2,3).

These findings indicate that our chromatin loop identification method and analysis pipeline effectively establish connections between chromatin loops and gene functions.

### 5. The signature of single-cell chromatin loops refines cell subtypes

To further assess CellLoops capability in refining cell states, we applied it to dual-modality data from mouse embryonic development obtained using HiRES technology at 100Kb resolution. In a previous study, these single cells were well-annotated based on RNA sequencing data using specifically expressed marker genes. Compared to Higashi, which utilizes single-modality 3D genome maps, CellLoops detection of single-cell chromatin loops delineated cell types more distinctly by mitigating cell cycle effects (Fig. 5a). Additionally, when compared to single-cell gene expression features, CellLoop revealed finer sub-clusters within the same cell type, indicating more detailed cell subtypes (Extended Data Fig. 5a, 6a–c).

**Fig. 5:**
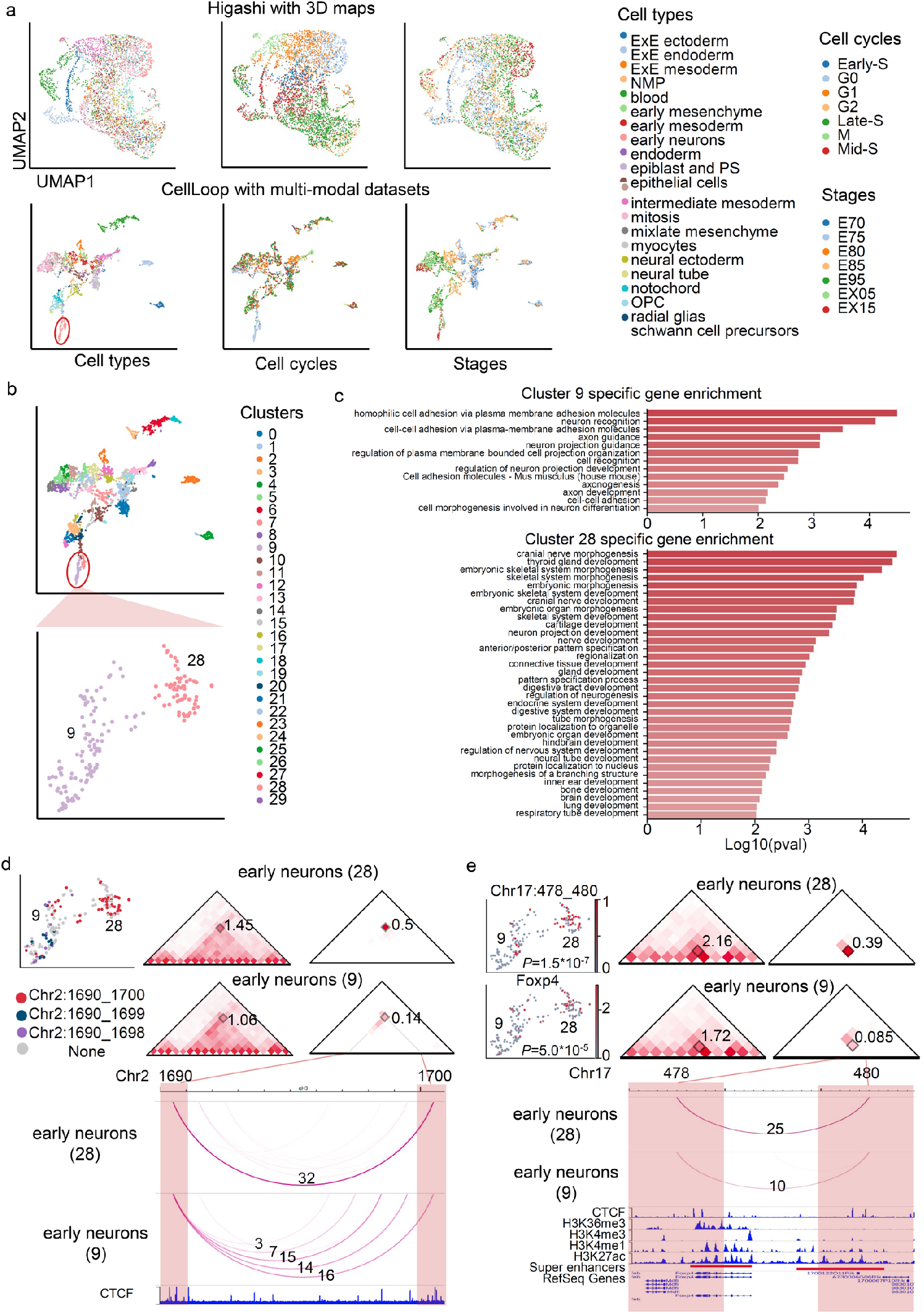
Application of CellLoop to the HiRES Dataset for Refining Cell Subtypes. **a**, UMAP visualization of cells using cell embeddings from the Higashi method (top) and single-cell chromatin loop features (bottom). Cells are labeled by cell type, cell cycle, and cell stage from left to right. **b**, Upper: UMAP visualization of cells using single-cell chromatin loop features, with cells labeled by clustering via the Leiden algorithm (resolution = 5). Bottom: UMAP visualization of clusters 9 and 28. **c**, Gene enrichment analysis for two gene sets associated with loop-gene pairs in clusters 9 and 28. These two cell clusters serve as reference sets for identifying loop-gene pairs. **d**, Example of a specific chromatin loop (Chr2:1690_1700) in cluster 28. Upper left: UMAP visualization of the Chr2:1690_1698/1699/1700 chromatin loop across cells. Upper middle and right: CFmap and LFmap surrounding the specific chromatin loop in the two clusters. Bottom: WashU Epigenome Browser showing the CTCF signal and LFmap for both clusters around the chromatin loop. Small squares mark the chromatin loops of interest. **e**, Example of a loop-gene pair in cluster 28. Upper left: UMAP visualization of the chromatin loop and corresponding gene expression. Upper middle and right: CFmap and LFmap surrounding the chromatin loop in the two clusters. Bottom: WashU Epigenome Browser displaying epigenetic signals and LFmap around the chromatin loop.

Notably, within the early neuron population, CellLoop identified two novel cell subtypes, designated as subtypes 9 and 28, based on UMAP clustering of single-cell chromatin loop features (Fig. 5c). A comparative analysis between these subtypes uncovered 254 and 420 chromatin loops specific to subtypes 9 and 28, respectively (Extended Data Fig. 7a, b; Supplementary Fig. 4, 5). For instance, a prominent chromatin loop (Chr2:1690_1700) was observed in subtype 28, whereas the corresponding region in subtype 9 exhibited a stripe architecture (Chr2:1690_1697/1698/1699/1700) (Fig. 5e), a structure consistent with the loop extrusion model facilitated by CTCF and Cohesin binding. Another example includes the formation of a significant chromatin loop (Chr18:763_768) in subtype 9, while the adjacent chromatin loop (Chr18:763_769) was notably present in both subtypes 9 and 28 (Extended Data Fig. 8a).

To explore potential associations between chromatin loops and gene expression in these subtypes, we identified 41 and 49 chromatin loop-specific gene expression pairs (loop-gene pairs) unique to subtypes 9 and 28, respectively (Extended Data Fig. 9a, b; Supplementary Fig. 6, 7). Gene enrichment analysis revealed that genes associated with subtype 9s loop-gene pairs were enriched in Gene Ontology terms related to cell adhesion, neuron recognition, and axon guidance (Fig. 5d, top). In contrast, genes linked to subtype 28s loop-gene pairs were associated with terms pertinent to organ, tissue, and system development, as well as the regulation of developmental processes (Fig. 5d, bottom). These loop-gene pairs frequently connected promoters with distal regulatory elements. Specifically, in subtype 28, the chromatin loop Chr17:478_480 likely brings a super-enhancer into proximity with the Foxp4 gene within the nuclear space, potentially activating Foxp4 expression (Fig. 5f). Previous studies have shown that Foxp4 is primarily expressed in lung, neural, and intestinal tissues during embryonic development^23^. In subtype 9, the chromatin loop Chr16:423_427 may regulate the Gap43 gene by linking it to a distal enhancer (Extended Data Fig. 8b); Gap43 is known to be essential for various aspects of mouse embryonic development, particularly in neuronal growth, synaptic plasticity, and cell migration^24^.

**Fig. 6:**
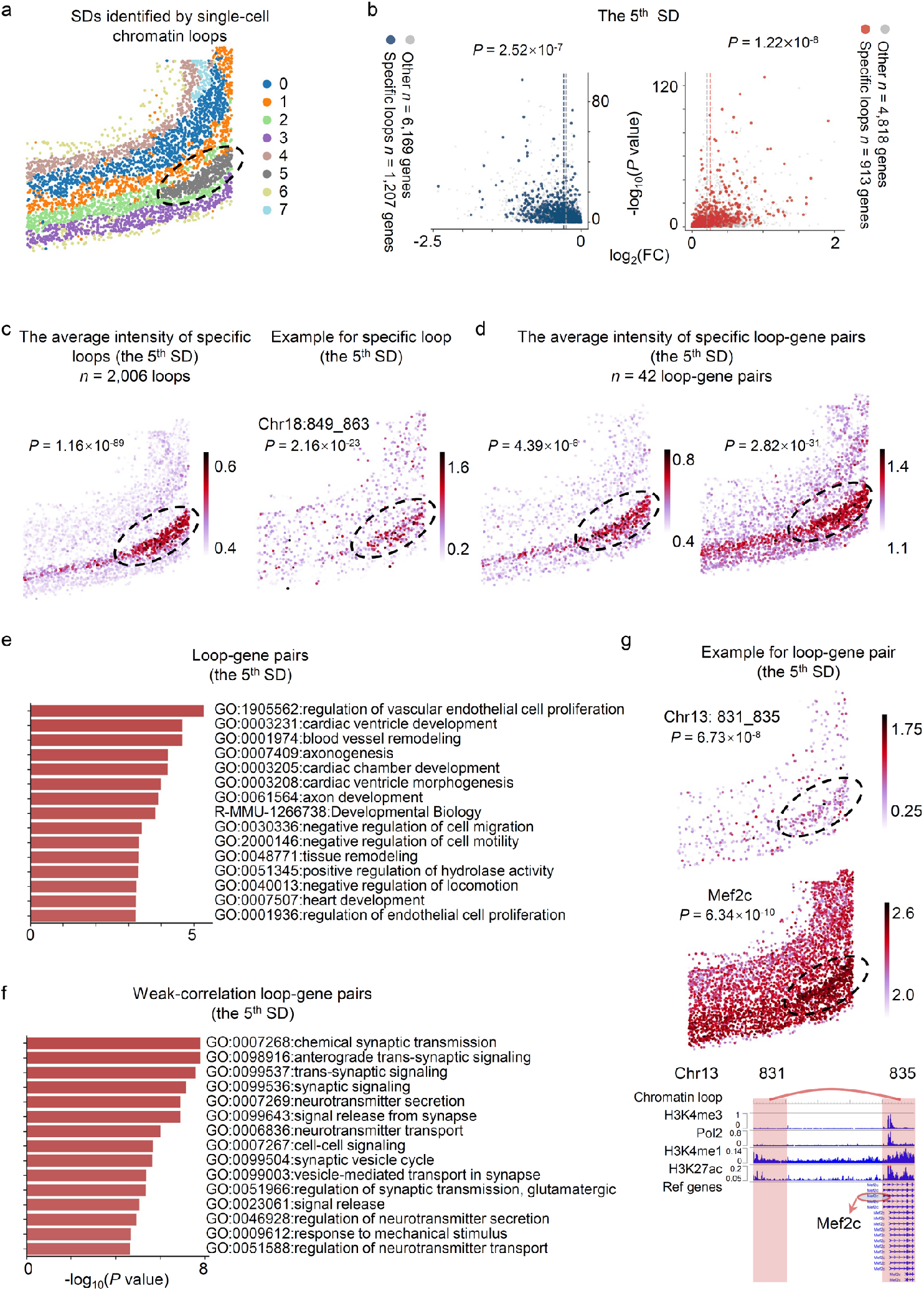
Application of CellLoop to the GAGE-seq and MERFISH Datasets for Depicting Spatial Domains. **a**, Spatial domains (SDs) identified by single-cell chromatin loops. The 5^th^ SD is highlighted with an ellipse. **b**, Left: Volcano plot of differential expression. Blue circles denote low-expressed genes within the anchor regions of chromatin loops specific to the 5^th^ SD, compared to other cells. Grey circles represent other low-expressed genes. Right: Red circles denote high-expressed genes within the anchor regions of chromatin loops specific to the 5^th^ SD, while grey circles represent other high-expressed genes (P values derived from one-sided *t*-tests). **c**, Left, in situ plot of the average intensity of the 5^th^ SD specific chromatin loops. Right, in situ plot of the example chromatin loop Chr18:849_863. **d**, In situ plots of the average intensity of loop-gene pairs specific to the 5^th^ SD (left: chromatin loops; right: genes). **e**, Gene enrichment analysis of the gene sets associated with loop-gene pairs in the 5^th^ SD. **f**, Gene enrichment analysis of the gene sets associated with weak-correlation loop-gene pairs in the 5^th^ SD. **g**, Upper and middle, In situ plots of example loop-gene pair. Bottom, WashU Epigenome Browse shows epigenetic signals around the chromatin loop.

In summary, CellLoop enhances the understanding of subtle biological differences among single cells from the perspective of chromatin loops, enabling a more detailed characterization of finer cell subtypes.

### 6. Single-cell chromatin loop signatures enhance spatial domain analysis

Recent advancements have enabled simultaneous mapping of single-cell 3D genomes and transcriptomes in the mouse cortex using GAGE-seq^6^. Additionally, MERFISH (multiplexed error-robust fluorescence in situ hybridization) has effectively identified transcriptomes and spatial locations of single cells in the mouse primary motor cortex^25^. To assess CellLoops capability in resolving spatial domains (SDs) from a 3D genome perspective, we applied it to the GAGE-seq dataset to identify single-cell chromatin loops at 100Kb resolution. Utilizing the transcriptomic modality of GAGE-seq as an intermediary, we established a connection between the 3D genome and spatial locations using Tangram^26^, thereby revealing spatial patterns of single-cell chromatin loops (Methods).

We then employed single-cell chromatin loops as characteristic features to identify SDs via STAGATE^27^ (Fig. 6a). The alignment between the identified SDs and major cell types suggests an association between chromatin loops and cell types (Extended Data Fig. 10a). Focusing on the distinctive fifth SD, we categorized genes associated with chromatin loops into high-expression and low-expression groups. Within this SD, 2,120 specific chromatin loops corresponded to significantly higher or lower expression levels (*t*-test *P*=1.22×10^−8^ for the higher expression group and *P*=2.52×10^−7^ for lower expression group, Fig. 6b). A similar analysis of the second SD also demonstrated a clear association between differentially expressed chromatin loops and differential gene expression patterns (*t*-test *P*=1.88×10^−24^ for higher expression and *P*=6.85×10^−6^ for lower expression, Extended Data Fig. 10b).

Further analysis of the fifth SD, using the second SD as a reference, identified 2,006 chromatin loops (*t*-test *P*=1.16×10^−89^, Fig. 6c, and Extended Data Fig. 10c), along with 42 loop-gene pairs showing significant correlations between specific chromatin loops and specifically expressed genes (*t*-test for loops *P*=4.39×10^−6^ and for genes *P*=2.82×10^−31^, Fig. 6d), all specific to the fifth SD. We designated this collection as the gene set propelled by chromatin loops (CL-genes). In contrast, the gene set driven by chromatin loops within the fifth SD encompassed a broader spectrum of biological functions compared to that of the second SD (Fig. 6e, Extended Data Fig. 10d, and Supplementary Tables 1 and 2).

To further dissect the functional uniqueness of CL-genes, we identified 29 weak-correlation loop–gene pairs within the fifth SD, defined as normally specifically expressed genes not driven by chromatin loops (normal-genes). Enrichment analysis revealed that CL-genes are associated with terms such as axonogenesis, axon development, cell migration, cell motility, locomotion, and proliferation, whereas normal-genes are linked to synaptic signaling, transmission, glutamatergic pathways, cell-cell signaling, signal release, neurotransmitter secretion, and transport (Fig. 6e,f and Supplementary Tables 1 and 3). The significant functional differences between the two gene sets suggest that genes driven by chromatin loops possess distinct functional specificity.

Focusing on specific gene loci driven by chromatin loops in the fifth SD, previous studies have confirmed that abnormal expression of the Mef2c gene is closely associated with autism spectrum disorder. Moreover, single-base editing technology has been successfully used to improve neurodevelopment and core autistic-like behavioral phenotypes in Mef2c autistic mouse models^28^. In our study, we found that the Mef2c gene is likely regulated by an enhancer, a distal regulatory element at a locus (83.1-83.2 Mb) with significant enhancer signal enrichment, resulting in increased expression levels (Fig. 6g). Additionally, Hbegf has been found to act as a mediator for cancer cell passage through the blood-brain barrier, enhancing cancer cell migration and invasion into brain tissues^29^. We observed that Hbegf expression might be regulated by a distal enhancer. Similarly, the EphA4 gene, implicated in neurodegenerative diseases such as amyotrophic lateral sclerosis, exhibits a comparable distal regulatory pattern^21, 30^.

Overall, CellLoop effectively and accurately identifies long-range regulatory relationships, which may play a crucial role in maintaining normal biological functions.

## Discussion

CellLoop accurately identifies chromatin loops in individual cells by formulating the task as a density-based peak detection problem, considering the multi-level background strength of chromatin interactions^16^. Utilizing two straightforward metrics in parallel, CellLoop efficiently detects chromatin loops across tens of thousands of cells within a reasonable timeframe. The resulting LFmap offers enhanced biological interpretability compared to the raw CFmap. Furthermore, we have developed a computational pipeline to elucidate the functional implications of single-cell chromatin loop features identified by CellLoop. Applying CellLoop to various single-cell 3D genomic datasets, we demonstrate its capability to extract unique chromatin loop signatures, facilitating the discovery of subtle cellular biological differences and enabling the reinterpretation of cell states or spatial domain functions through 3D genomic chromatin loops.

CellLoop employs an unsupervised approach to distinguish chromatin loops from existing chromatin interactions within the data. Future directions include constructing realistic nuclear architectures by integrating multi-modal information to compensate for missing signals, thereby enabling the identification of de novo multi-way chromatin loops. Another limitation of CellLoop is its reliance on the specific biological context of single cells. Therefore, developing computational models that integrate multi-omics, multi-modalities, and imaging data^31, 32^ is essential to elucidate intercellular associations and establish a robust biological context for single cells. Additionally, with the rapid advancement of spatial 3D genomics data^33^, identifying spatially associated chromatin loops will further enhance the analysis of spatial domains and cell-to-cell communication from the perspective of 3D chromatin loops. Finally, a significant future direction involves understanding how alterations in 3D structural patterns impact biological processes and the onset and progression of diseases by exploring the link between multi-scale 3D structural patterns and DNA modifications^34^.

In summary, with the rapidly increasing volume of single-cell 3D genomic data, CellLoop is poised to play a pivotal role in rapidly and accurately identifying chromatin loops at single-cell resolution in the context of diseases or specific biological states. The unique attributes characterized by single-cell chromatin loops will empower precise refinement of cell annotations, facilitate comprehensive characterization of gene regulatory networks, and drive in-depth reanalysis of cell functions.

## Methods

### Identification of single-cell chromatin loops

The CellLoop algorithm comprises two primary components: the preference prediction module and the peak calling module (Fig. 1a). The algorithm inputs include raw single-cell contact maps and initial embedding matrices (Methods). Focusing on intra-chromatin interactions, we illustrate the algorithm using a specific chromatin region from single cell *v* as an example.

In the first component, we constructed an undirected graph *A*^*v*^ ∈ {0,1}^*n*×*n*^ with bins as nodes, derived from the raw contact map of single cell *v* (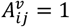 if there are interactions bin *i* and bin *j*, otherwise, 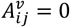). Additionally, we generated a weighted undirected k-nearest neighbors (*k*-NNs) graph *W*^*v*^ ∈ *R*^*m*×*m*^ with cells as nodes, connecting single cell *v* to its *k*-NNs based on Euclidean distances calculated from the initial embedding space (Fig. 1a, top). The enhanced contact map *S*^*v*^ for single cell *v* is defined as:

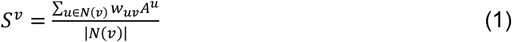

where *N*(*v*) represents neighboring cells of single cell *v* with weights *w*_*uv*_ > 0.2. We discuss the selection of the default *k* value and its impact on algorithm robustness in the Supplementary Notes. Next, we predicted the preference for chromatin interaction formation *ρ* by integrating interaction strengths from multiple levels (Fig. 1a, bottom), defined as:

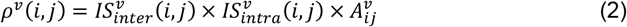

where 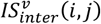 and 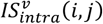 represent inter-cellular and intra-cellular interaction strengths between bin *i* and bin *j*, respectively. 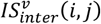 is defined as:

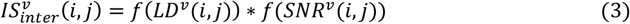

where *LD*^*v*^(*i, j*) denotes the local strength of chromatin interactions (*i, j*) from neighboring inter-cellular maps:

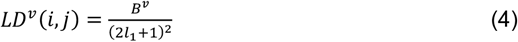

where 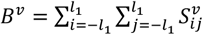 (default *l*_1_ = 1; see Supplementary Fig. 8). *SNR*^*v*^(*i, j*) is defined as

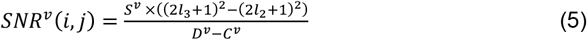

where 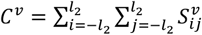 and 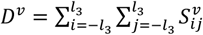 (default *l*_2_ = 2, *l*_3_ = 5; see Supplementary Fig. 8). The function *f* normalizes these values, defined as 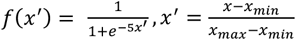. The intra-cellular interaction strength 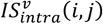 is defined as:

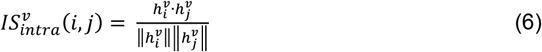

where 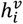 represents the embedding vector of bin *i*, calculated by applying the node2vec graph representation learning algorithm to the undirected graph *A*^*v*^ based on a biased random walk procedure^35^.

The second component of the CellLoop algorithm distinguishes chromatin loops from the preference map *ρ* by formulating the task as a two-dimensional peak calling problem. We retained chromatin interactions with *ρ* > 0 as candidates for peak calling and defined another metric—the minimum distance *δ*^*v*^(*i, j*) to higher-strength chromatin interactions within a range of *Dis*_*max*_ (default *Dis*_*max*_ = 100):

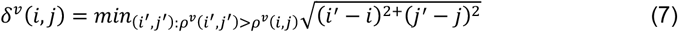

where the maximum value of *δ*^*v*^(*i, j*) is less than *Dis*_*max*_. The overall score for chromatin interaction (*i, j*) is then:

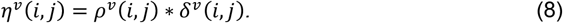

We ranked candidate chromatin interactions by *η*^*v*^ in descending order, and Both *η*^*v*^ and their rank *r*^*v*^ are normalized by 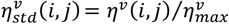 and 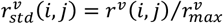, respectively. To distinguish true chromatin loops from other interactions, we identify the inflection point *i*_*bp*_, *j*_*bp*_ by minimizing the sum of squares of the normalized scores:

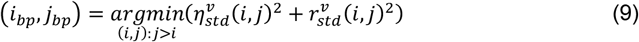

Finally, the set of chromatin loops is extracted by selecting interactions with 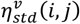 greater than 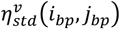:

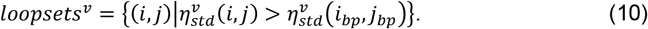

### The pipeline for dividing single-cell chromatin loops into different types

To facilitate integrated analysis across multiple cell types, we developed a computational pipeline to classify detected chromatin loops into distinct subtypes (Extended Data Fig. 2). The pipeline inputs include the single-cell chromatin loops matrix, *M* ∈ {0,1}^*n*×*m*^ and cell type annotation vector of single cells *V* ∈ {1,2, …, *N*}^*n*^, where *m* represents the number of cells and *m* denotes the number of chromatin loops. *M* indicates the presence (1) or absence (0) of specific chromatin loops in individual cells, while *V* provides the corresponding cell type annotations. Prior to analysis, we implemented a quality control filter, retaining only chromatin loops detected in at least 10 single cells. For a given chromatin loop *i* and cell type *j*, we defined the positive sample set (Pos) as the occurrence vector of loop *i* in cells annotated as type *j*. The negative sample set (Neg) was constructed by performing multiple downsamplings (default: 10 iterations) of the occurrence vector of loop *i* in cells annotated as other cell types, ensuring that |Neg| = |Pos|. We then calculated the average enrichment score (ES) and statistical significance (*P* value) of chromatin loops in Pos compared to Neg. The ES was defined as the ratio of the proportion of loop occurrences in Pos to that in Neg. *P* values were computed using a t-test and adjusted for multiple testing using the Benjamini-Hochberg method^36^. Based on these metrics, chromatin loop *i* was categorized into one of four specificity levels for cell type *j*, ordered from highest to lowest specificity: strong specific chromatin loop (strong-loop), weak specific chromatin loop (weak-loop), no specific chromatin loop (noSpe-loop), and disappearing chromatin loop (Dis-loop) (Extended Data Fig. 2). Furthermore, considering the distribution of specificity levels of loop *i* across different cell types, each loop was classified into one of three categories: specific to an individual cell type (Indi-Spe), specific to multiple cell types (Multi-Spe), or common (Com) (Extended Data Fig. 2). Focusing on Indi-Spe chromatin loops, we further classified them based on the gene expression status of their anchor regions relative to other cell types into three categories: gene specifically highly expressed (Indi-Spe-highgene), gene specifically lowly expressed (Indi-Spe-lowgene), and gene non-specifically expressed (Indi-Spe-nonSpegene). In this context, we selected the most specific gene within the anchor regions of each chromatin loop for analysis.

For analyses targeting specific cell types or subtypes, we employed the Scanpy package to identify specific chromatin loops and genes by specifying the reference cell set for comparison using a one-sided *t*-test^39^.

### The definition of loop-gene pairs

In single-modality analyses, we define a *loop-gene pair* as a chromatin loop that exhibits specificity and encompasses within its anchor regions a gene that is highly and specifically expressed (selecting the most specific gene). For dual-modality data, an additional criterion is applied: the Pearson correlation coefficient between the single-cell expression level of the gene and the normalized occurrence score of the chromatin loop, specific to the pertinent cell type or subtype, must exceed 0.1. Pairs exhibiting a correlation coefficient of 0.1 or below are categorized as *weak-correlation loop-gene pairs*.

### The initial embedding matrices of single cells

The CellLoop algorithm initializes cell embeddings through distinct approaches tailored to specific data modalities. **Single-modality 3D genomic data:** For datasets comprising solely 3D genomic information, CellLoop employs Higashi^12^, an integrative framework based on hypergraph representation learning, to derive initial cell embeddings. **Dual-modality data integrating 3D genomics and other omics:** In scenarios where 3D genomic data is complemented by additional omics layers, CellLoop utilizes UMAP to generate initial cell embeddings, leveraging the supplementary omics data (refer to Methods: UMAP Visualization and Clustering of Single Cells).

Looking ahead, advancements in spatial 3D genomics are anticipated to enable the construction of initial cell embeddings by incorporating spatial coordinates, thereby enhancing the contextual understanding of chromatin architecture^33^.

### Quantitative performance evaluation metrices

We employed two metrics—genome spatial distance (GSD) score and compartment property consistency (CPC) score—to quantitatively assess the performance of detected chromatin loops in comparison to control interactions. For each chromatin loop, control chromatin interactions were constructed by fixing one anchor of the loop and positioning the other anchor at the same distance in the opposite direction, equal to the length of the chromatin loop. These control interactions were required to be present in the raw single-cell contact map.

#### GSD score

High-quality 3D genome structures for single cells were generated at a 20KB resolution, as previously described^2, 37^. The GSD score for a chromatin loop was calculated by determining the Euclidean distance between the two anchors in 3D genomic space. A lower GSD score indicates higher-quality chromatin loops.

#### CPC score

The scA/B values were computed from raw contact maps of single cells using the “dip-c color2” function^2, 38^. The CPC score for a given single cell was defined as the Pearson correlation coefficient between the scA/B values at the two anchor regions of the chromatin loop identified within that cell. A higher CPC score reflects better-quality chromatin loops in the single-cell analysis.

### Aggerated peak score (APS)

We conducted the APS analysis on two matrices at a 20KB resolution: the loop frequency map (LFmap) and the contact frequency map (CFmap). To quantify the aggregate enrichment of a set of chromatin loops in these matrices, we calculated the average of a series of submatrices extracted from both maps. Each submatrix is a 220KB × 220KB square, centered on a single chromatin loop located in the upper triangle of the matrices. The APS of a set of chromatin loops was determined by the ratio of the central pixel value to the mean of the pixels in the lower-left corner of the submatrices that exceed the average value.

### The enrichment analysis of ChIP-seq peak on chromatin loops

We retrieved ChIP-seq peaks for various epigenetic factors from ChIP-Atlas (https://chip-atlas.org/)^39^. Enrichment analysis was performed at a 20KB resolution. To quantify the aggregate enrichment of a set of chromatin loops, we centered on each chromatin loop and computed the number of peaks associated with all chromatin interactions surrounding the loop. This yielded a peak enrichment submatrix for each chromatin loop. The final enrichment matrix was obtained by calculating the average of a series of submatrices derived from the set of chromatin loops. The matrix was visualized using the plot_surface function in the matplotlib package.

### UMAP visualization and clustering of single cells

UMAP visualization and clustering of single-cell chromatin loop and RNA-seq matrices were performed using the Scanpy package^37^. For the single-cell chromatin loop matrices, we first filtered out chromatin loop features at a 100KB resolution that were observed in fewer than 10 cells. The filtered matrix was then normalized using the TF-IDF transform algorithm as previously described^40^. A lower-dimensional representation of the data was generated by incorporating the first 20 dimensions from the singular value decomposition (SVD) of the TF-IDF-transformed matrix. This reduced representation was subsequently used as input for the UMAP algorithm. Cell clusters were identified in the two-dimensional space using the Leiden clustering algorithm with default parameters.

For the single-cell RNA-seq matrix, cells with fewer than 100 genes and genes expressed in fewer than 5 cells were excluded. The remaining count data were normalized by multiplying each cells library size by a scale factor (default value = 10,000). The normalized matrix was then log-transformed in preparation for downstream analysis. Principal component analysis (PCA) was performed to reduce the dimensionality, and the resulting representation was used as input for UMAP. The Leiden clustering algorithm with default parameters was then applied to detect cell clusters.

For the single-cell scA/B matrix, the visualization and clustering pipeline followed a similar procedure to that of the RNA-seq matrix, with the exception that filtering of cells and features was not performed.

### Inference chromatin loop features for MERFISH cells

Our analysis focused on excitatory neuron cells present in both the GAGE-seq and MERFISH datasets. To infer the intensity of chromatin loop features in the MERFISH spatial data, we employed the Tangram algorithm^26^. First, we selected 254 genes detected in the MERFISH data as training genes to identify the optimal alignment between the transcriptome libraries of GAGE-seq and MERFISH, using the cells mode. Subsequently, we used the project_cell_annotation function to transfer the chromatin loop features from GAGE-seq to the MERFISH dataset.

### Identification of spatial domains using chromatin loop features for MERFISH cells

Chromatin loop features of MERFISH cells were analyzed using the STAGATE package^27^ to identify spatial domains. Features detected in fewer than 50 cells were excluded from the analysis. The top 50 differentially expressed features for each cell type were then selected as input for the STAGATE algorithm. Count normalization for each cell was performed by scaling the library size with a factor of 10,000 (default value). The normalized matrix was subsequently log-transformed to prepare it for downstream analysis. To construct a cell spatial neighbor network, we set the rad_cutoff parameter to 60 and employed the train_STAGATE function to learn low-dimensional latent representations of cells. Finally, the Louvain algorithm was applied to cluster the spatial domains.

## Supporting information

Supplementary results, methods, figures, and table legends

## Dataset availability

In this work, we used several publicly available single-cell Hi-C datasets: Single-cell Dip-C for the developing mouse cortex and hippocampus^2^ (GSE146397), single-cell double-modality data of mouse embryonic development using HiRES^3^ (GSE223917), single cell double-modality data of mouse cortex using GAGE-seq^6^ (GSE238001). For all scHi-C datasets, we kept only the high-quality coverage cells with more than 200,000 read pairs. After filtering, the Dip-C dataset contains 1,812 cells, whereas the HiRES dataset contains 4,273 cells and the GAGE-seq dataset contains 1,925 cells. In single-cell maps of 20KB or higher resolution, CellLoop identifies chromatin loops within a 10Mb genomic distance; while in those of 10KB or lower resolution, it identifies such loops within 5Mb. The cell type annotation of Dip-C and HiRES datasets were provided in the original study, and cell types in GAGE-seq were determined using the CellTypist algorithm^41^ using single-cell RNA-seq for mouse brain cortex from Tasic et al as reference^42^. MERFISH spatial transcriptome dataset was downloaded from https://download.brainimagelibrary.org/cf/1c/cf1c1a431ef8d021/^32^. Single-cell RNA-seq datasets were downloaded from https://figshare.com/articles/dataset/sc_mouse_cortex/13677259/1?file=26404781.

## Code availability

The software package implementing the CellLoop algorithm has been deposited at Github (https://github.com/YusenYe/CellLoop).

## Acknowledgements

This work was supported by a grant from the Fundamental Research Funds for the Central Universities (no. QTZX23074 to Y.Y.), a grant from the Young Talent Fund of Association for Science and Technology in Xi’an (no. A023030001 to Y.Y.), National Natural Science Foundation of China grant (nos. 62132015, 62350087 and U22A2037 to L.G., no. 62002277 to Y.H., nos. 32341013, 12326614, 12126605 to S.Z., and Y.Y., and no. 62002275 to Y.Y.). We thank L. Zhang and W. Min for helpful discussions. We also thank the Key Laboratory of Computational Bioinformatics of Xi’an at Xidian University.

## Author contributions

Y.Y. conceived and designed the study. Y.Y. designed the CellLoop algorithm. Y.Y. implemented the CellLoop algorithm. L.G., H.C. and Y.H. provided additional input during the development of the methods. Y.Y. performed the data analysis. Y.Y. provided support for the development of the software packages. Y.Y. supervised the study. Y.Y. and S.Z. wrote the manuscript.

## Computing interests

The authors declare no competing interests.

---

**Extended Data Fig. 1:**
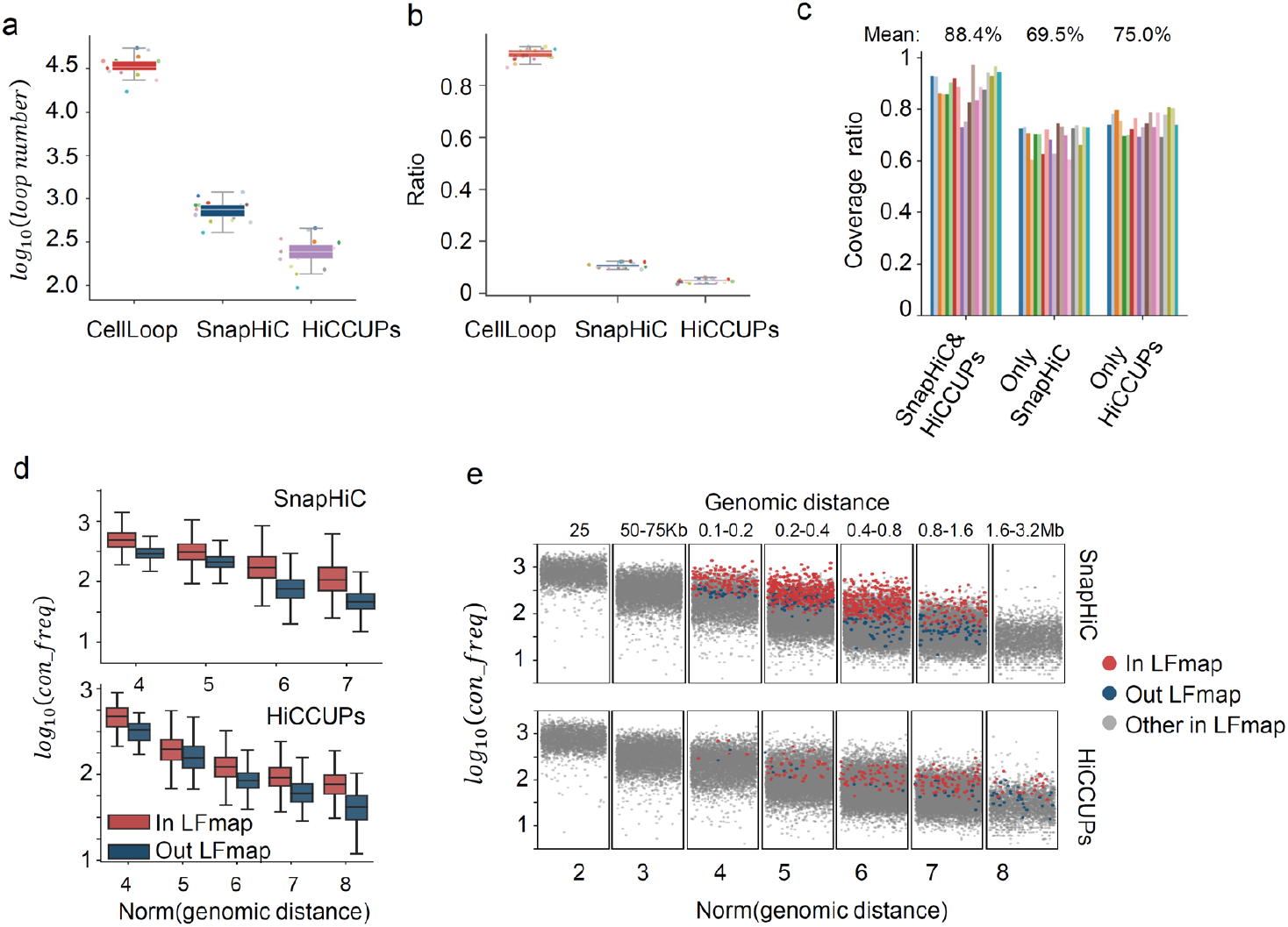
Comparison of LFmap with existing methods in the Dip-C dataset. **a**, The number of chromatin loops identified by Cellloop, HiCCUPs, and SnapHiC in different chromosomes. **b**, Genomic region coverage ratio of chromatin loops identified by CellLoop, HiCCUPs, and SnapHiC in different chromosomes. **c**, The ratio of chromatin loops detected by existing methods (SnapHiC&HiCCUPs CellLoop represents the chromatin loops detected simultaneously by the two methods) that are covered by LFmap. The average value of the coverage ratios for different chromosomes. **d**, The contact frequency distribution of chromatin loops detected by existing methods (SnapHiC and HiCCUPs). These loops are categorized into two groups based on their presence in LFmap. **e**, Scatter plot of the contact frequency of chromatin loops at different genomic distances. These loops are classified into three groups according to whether they are detected by existing methods (SnapHiC and HiCCUPs) or covered by LFmap.

**Extended Data Fig. 2:**
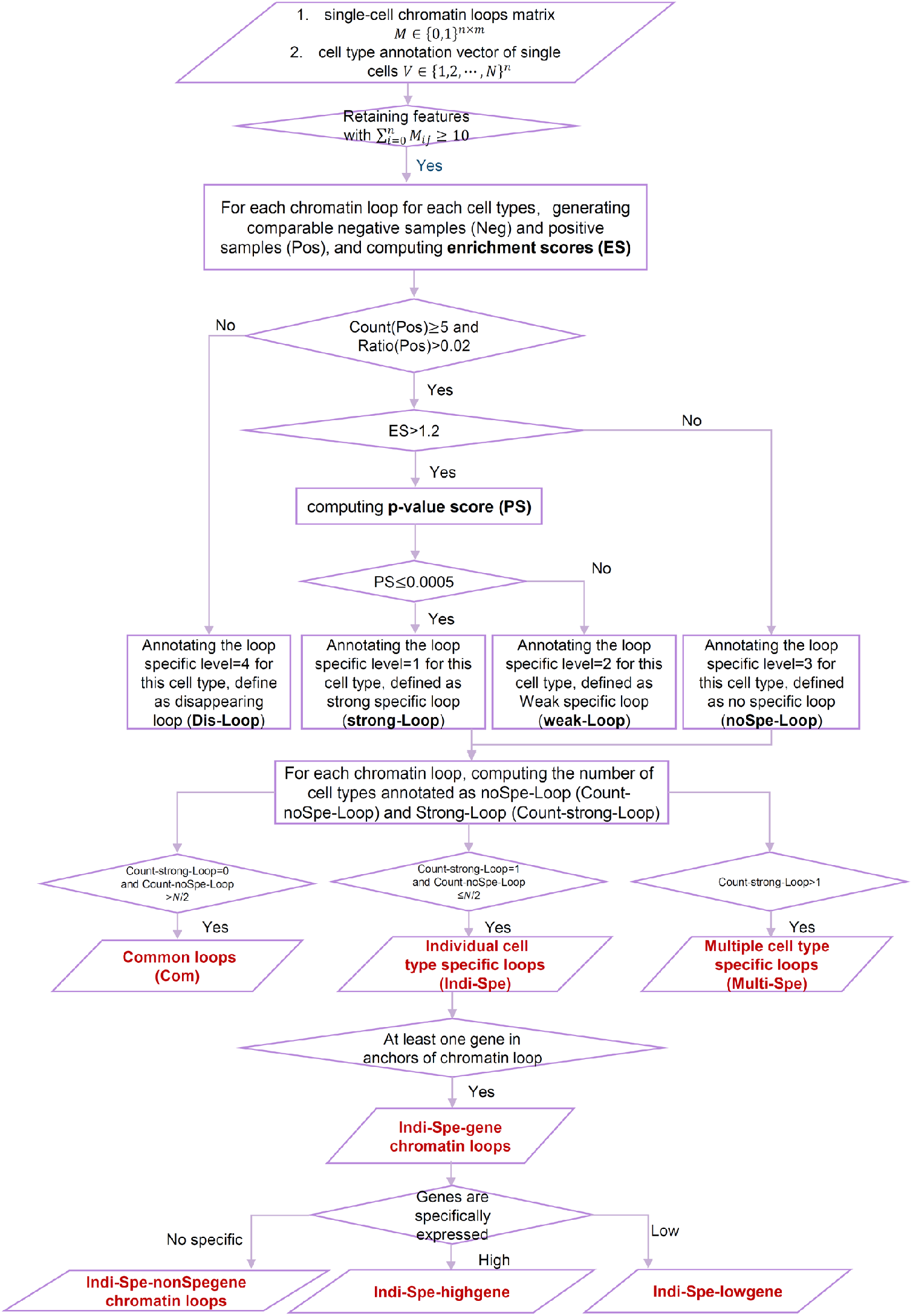
The pipeline for dividing single-cell chromatin loops into different types. The functions of Count(Pos) and Ratio(Pos) calculate the frequency and proportion of the given chromatin loop appearing in the Pos respectively. For detailed information, please refer to the Methods.

**Extended Data Fig. 3:**
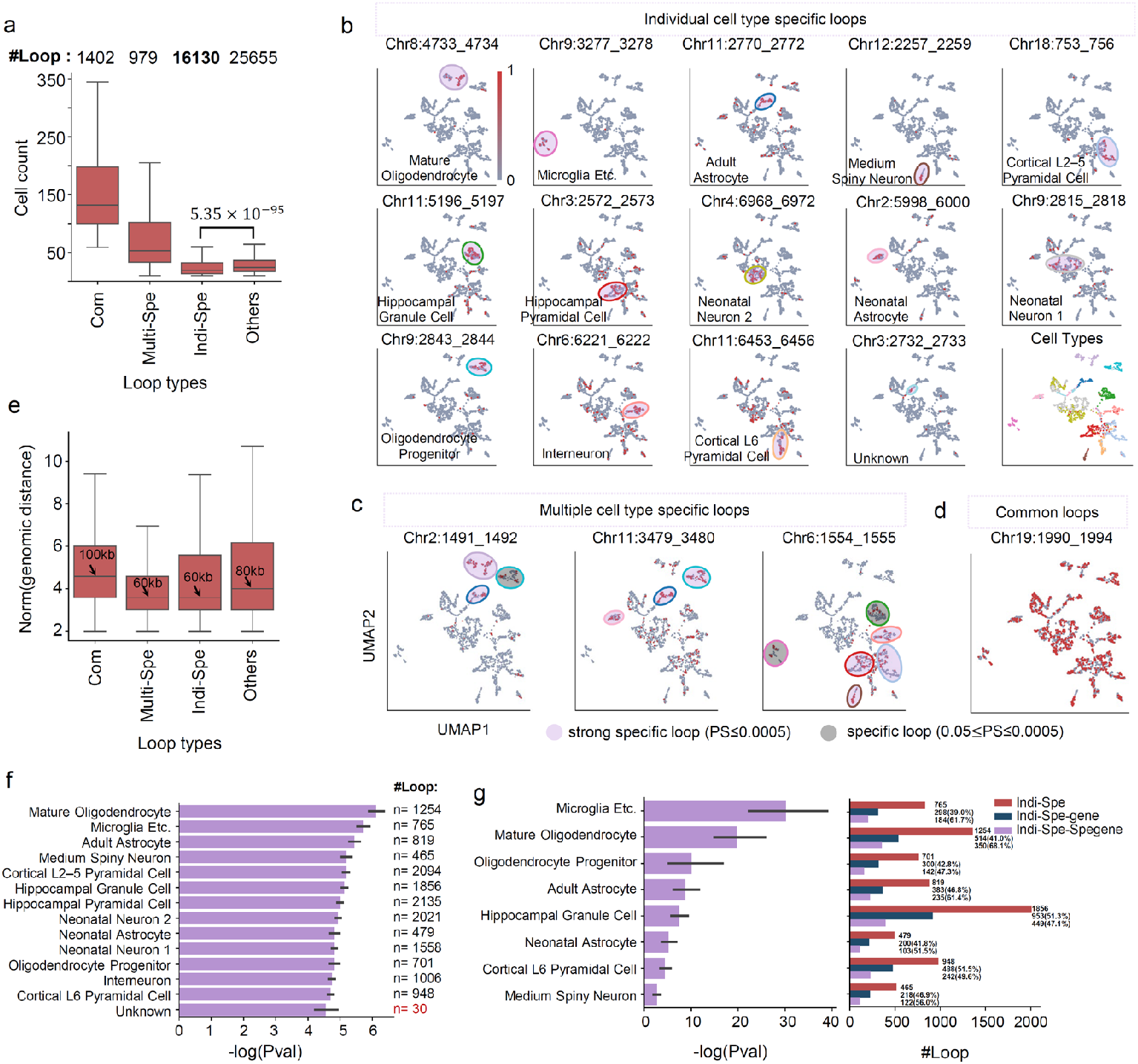
Statistics and visualization of different types of chromatin loops detected in the Dip-C data. **a**, Distribution of the number of cells detecting different chromatin loop types. The numbers in the upper of the figure represent the quantities of different chromatin loop types. **b**, UMAP visualization examples for Indi-Spe chromatin loops for each cell type. Ellipses mark the corresponding cell types. Lower right corner: UMAP visualization of cell types. **c**, UMAP visualization examples for Multi-Spe chromatin loops. Ellipses mark out corresponding cell types. **d**, UMAP visualization examples for Com chromatin loops. **f**, Significance score distribution and quantity of cell type differential chromatin loops. **g**, For eight one-to-one corresponding cell types between Dip-C and MALBAC-DT datasets, Left: significance score distribution of cell type differential chromatin loops, Right: Quantity and proportion of cell type differential chromatin loops under different conditions. Indi-Spe-Spegene includes two classes: Indi-Spe-highgene and Indi-Spe-highgene. See Extended Data Fig.2 for specific chromatin loop types.

**Extended Data Fig. 4:**
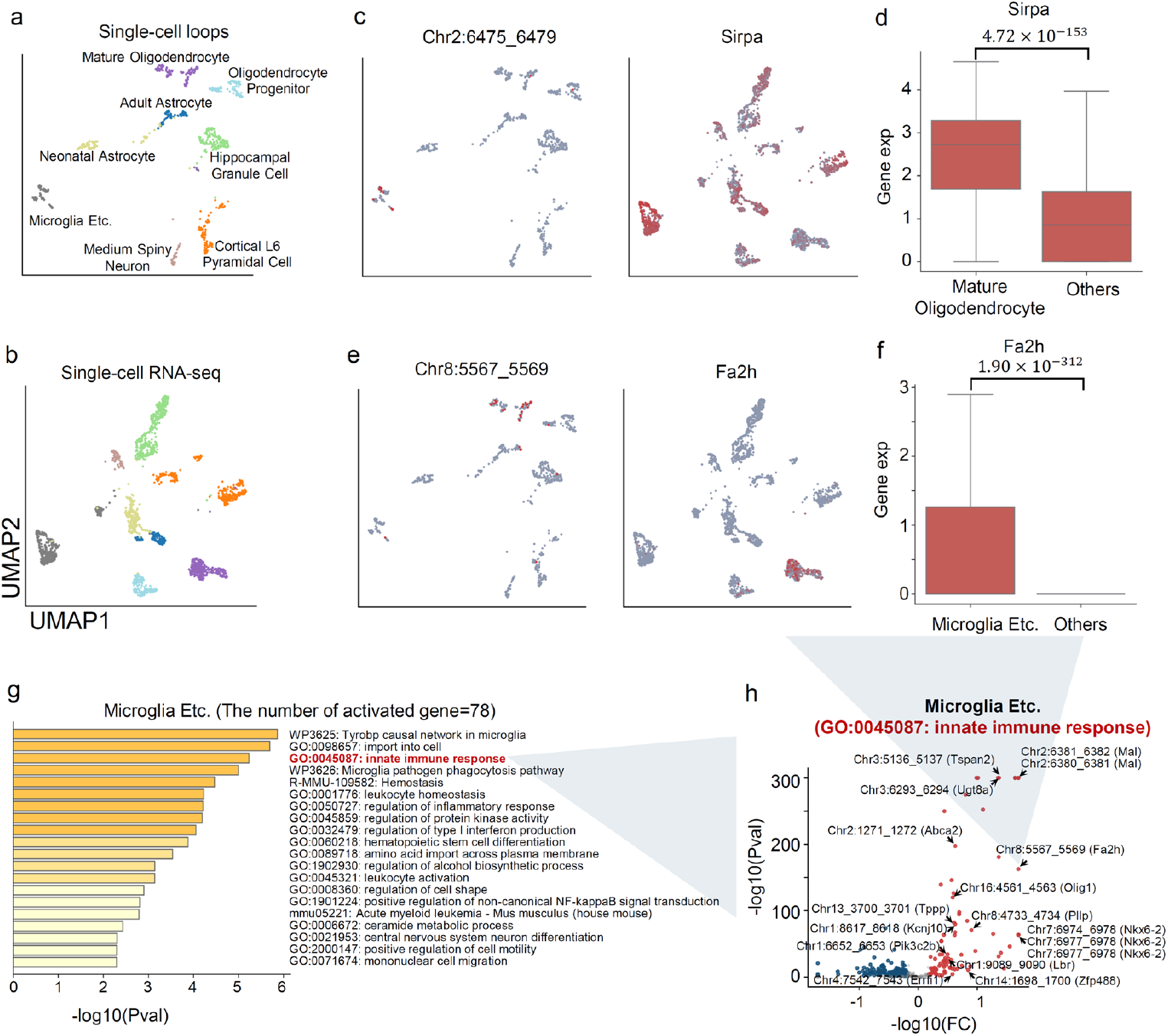
Analysis of the association between chromatin loops and gene functions for eight one-to-one corresponding cell types between the Dip-C and MALBAC-DT datasets. **a**, UMAP visualization using single-cell chromatin loop features. **b**, UMAP visualization using single-cell gene features. **c and e**, UMAP visualization of the occurrence of chromatin loops and the expression of their corresponding genes, respectively. **d and f**, Boxplot of the expression of genes in c and e, respectively. **g**, Gene function enrichment analysis for gene sets from Indi-Spe-highgene of Microglia Etc.. **h**, Volcano plot of Indi-Spe-highgene pairs for GO:0045087 term in Microglia Etc..

**Extended Data Fig. 5:**
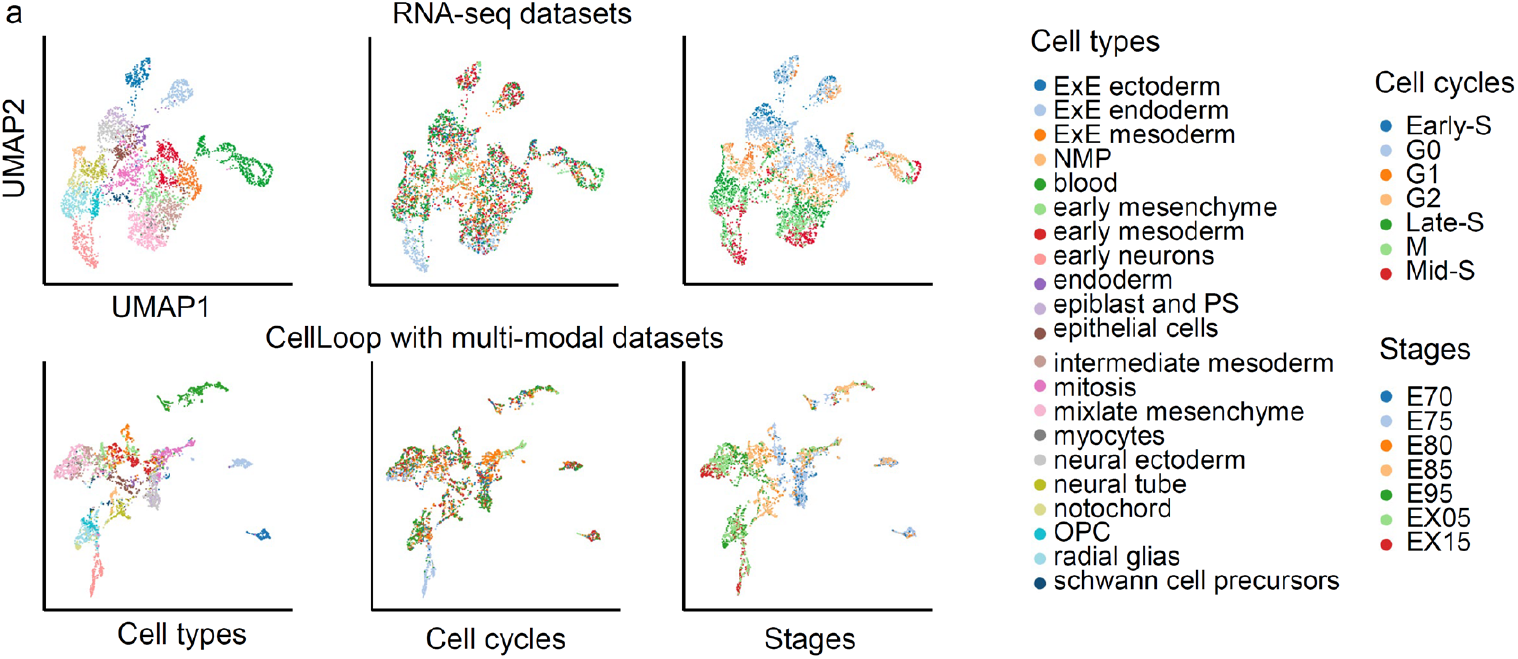
UMAP visualization of cells using gene feature from RNA-seq dataset(upper) and single-cell chromatin loops (bottom) features. Cells are labeled respectively by cell types, cell cycle, and cell stages from left to right, respectively.

**Extended Data Fig. 6:**
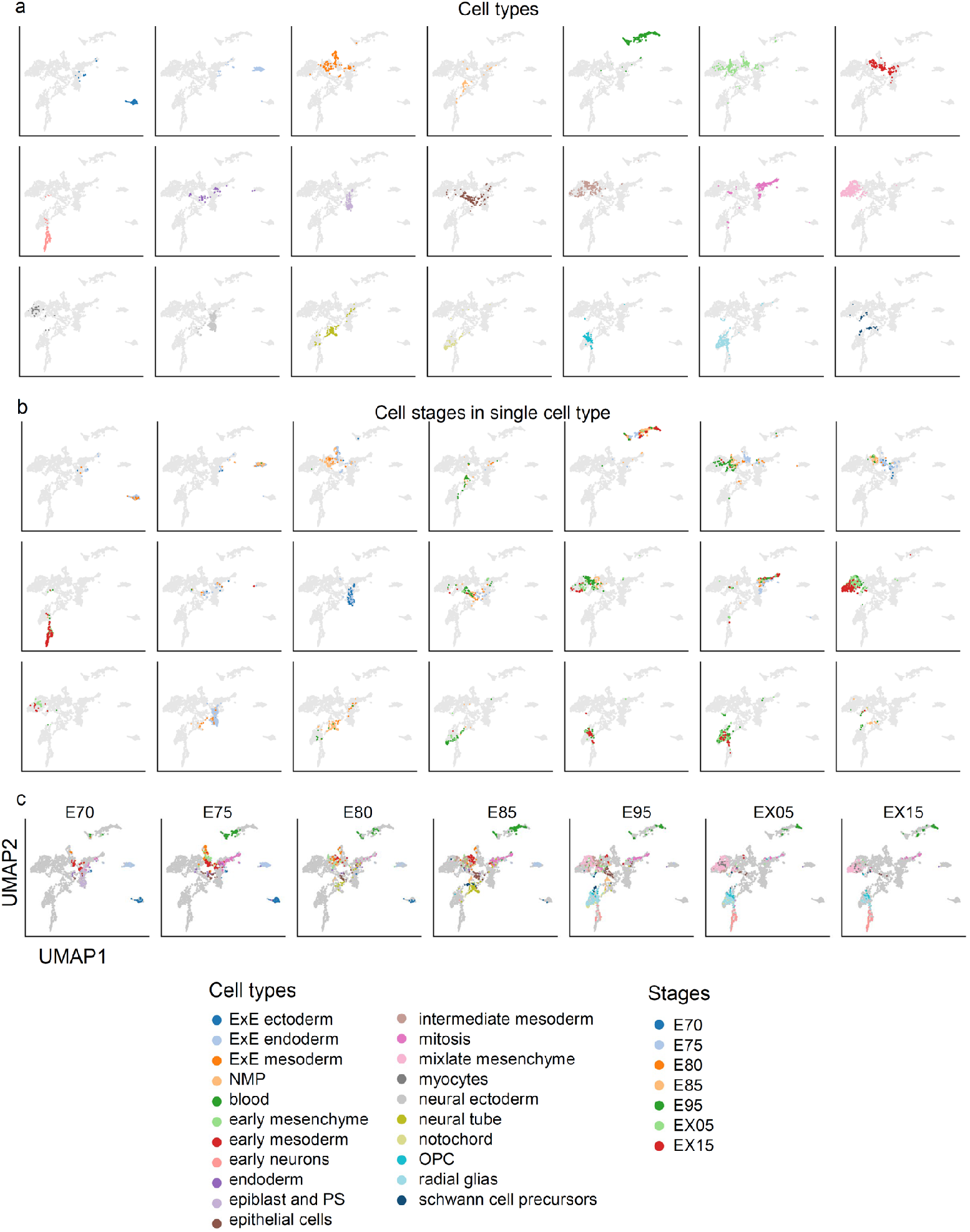
UMAP visualization of cells using single-cell chromatin loop features. **a**, Cells in each graph are labeled by one specific cell type. **b**, Each graph shows cells at cell stages of the same cell type. **c**, Each graph shows cell types in the same cell stage.

**Extended Data Fig. 7:**
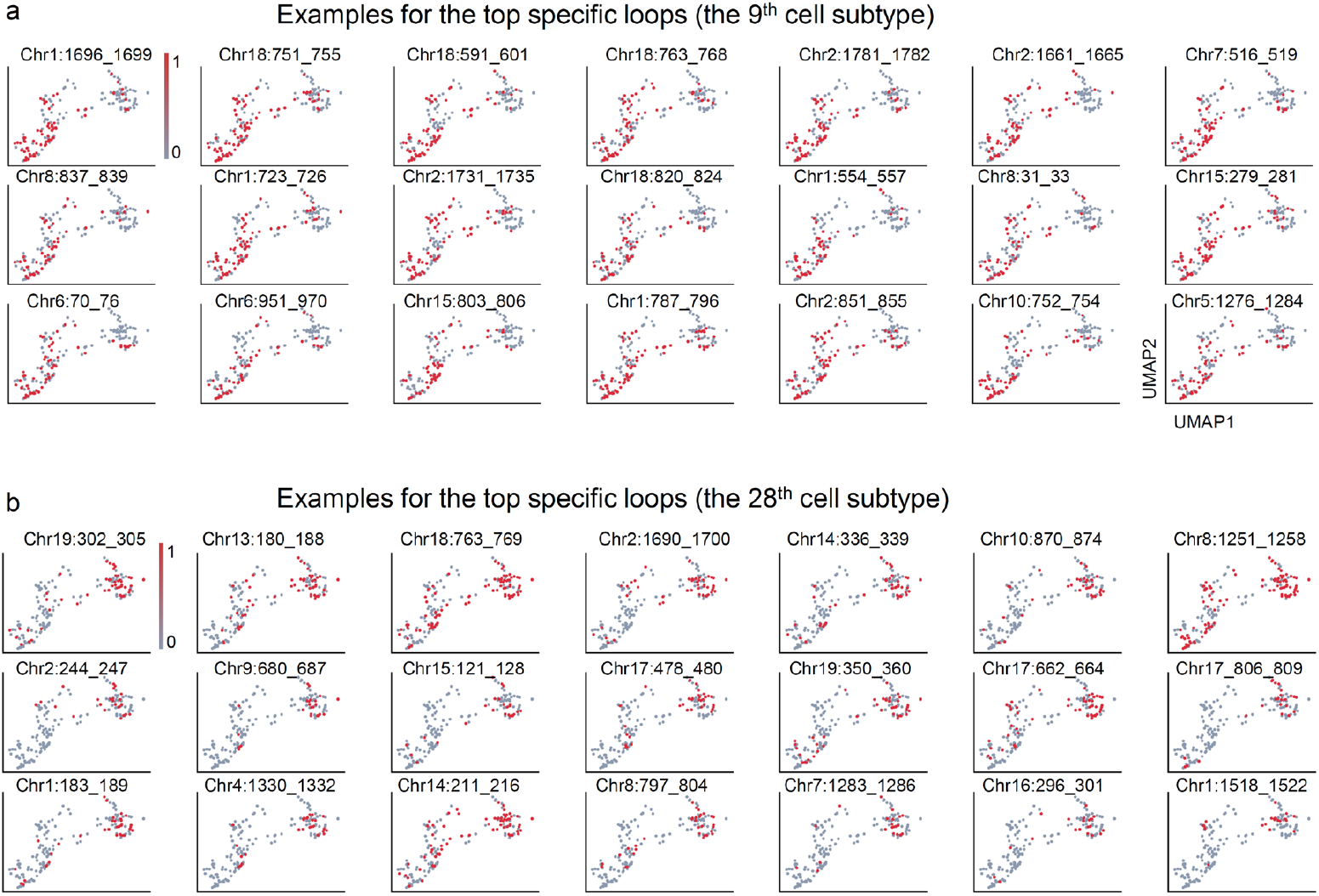
UMAP visualization of the top specific chromatin loops for clusters 9 (a) and 28 (b).

**Extended Data Fig. 8:**
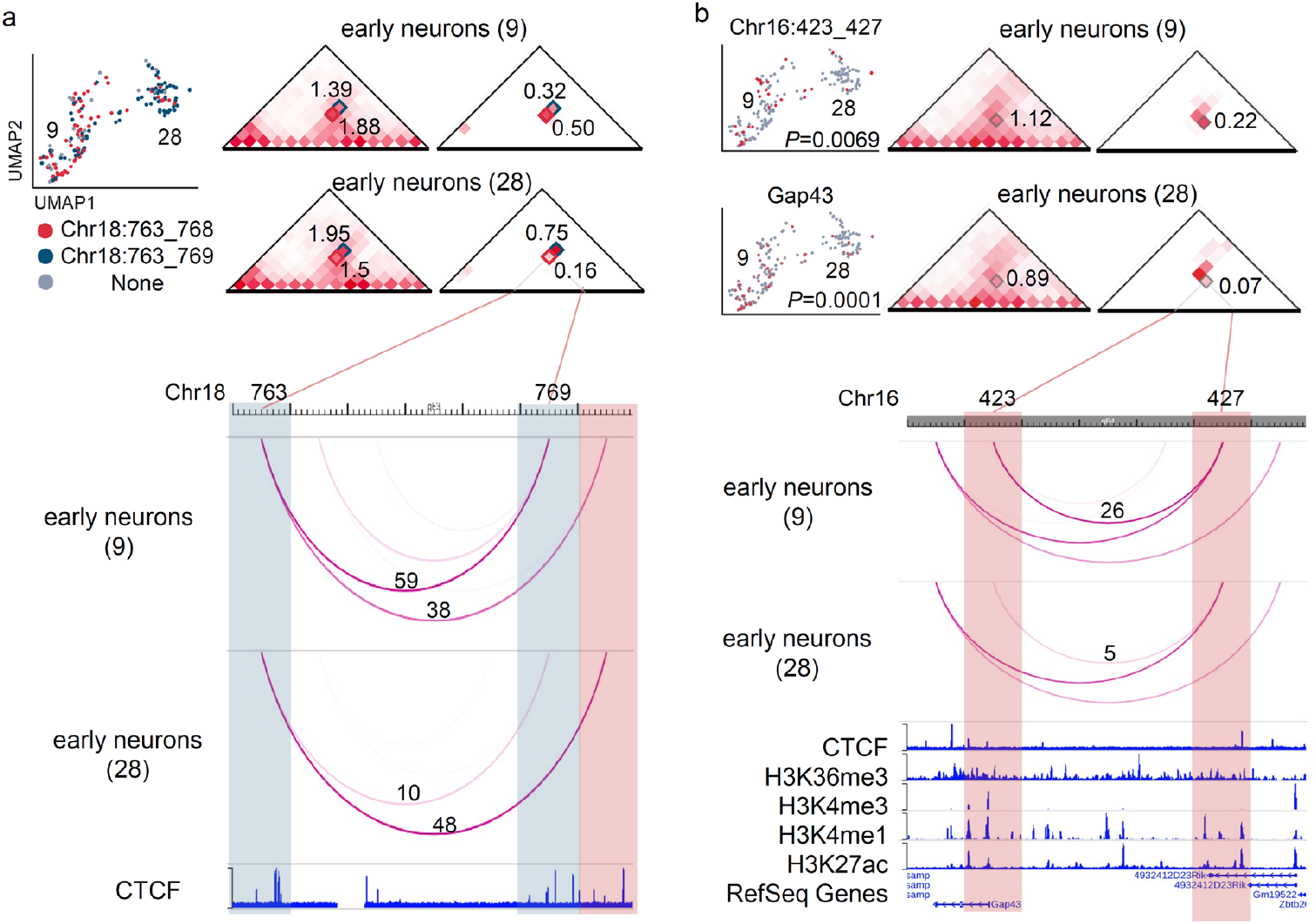
Examples for specific chromatin loop and loop-gene pair. **a**, An example of the specific chromatin loop Chr18:763_768 in cluster 9. Upper left, UMAP visualization of the Chr18:763_768/769 chromatin loop in cells. Upper middle and right, CFmap and LFmap around the specific chromatin loop in two clusters. Bottom, WashU Epigenome Browse shows the CTCF signal and LFmap of two clusters around the specific chromatin loop. Small squares mark the chromatin loops of interest. **b**, An example of loop-gene pair in cluster 9. Upper left, UMAP visualization of the chromatin loop and corresponding gene expression. Upper middle and right, CFmap and LFmap around the chromatin loop in two clusters. Bottom, WashU Epigenome Browse shows epigenetic signals and LFmap around the chromatin loop.

**Extended Data Fig. 9:**
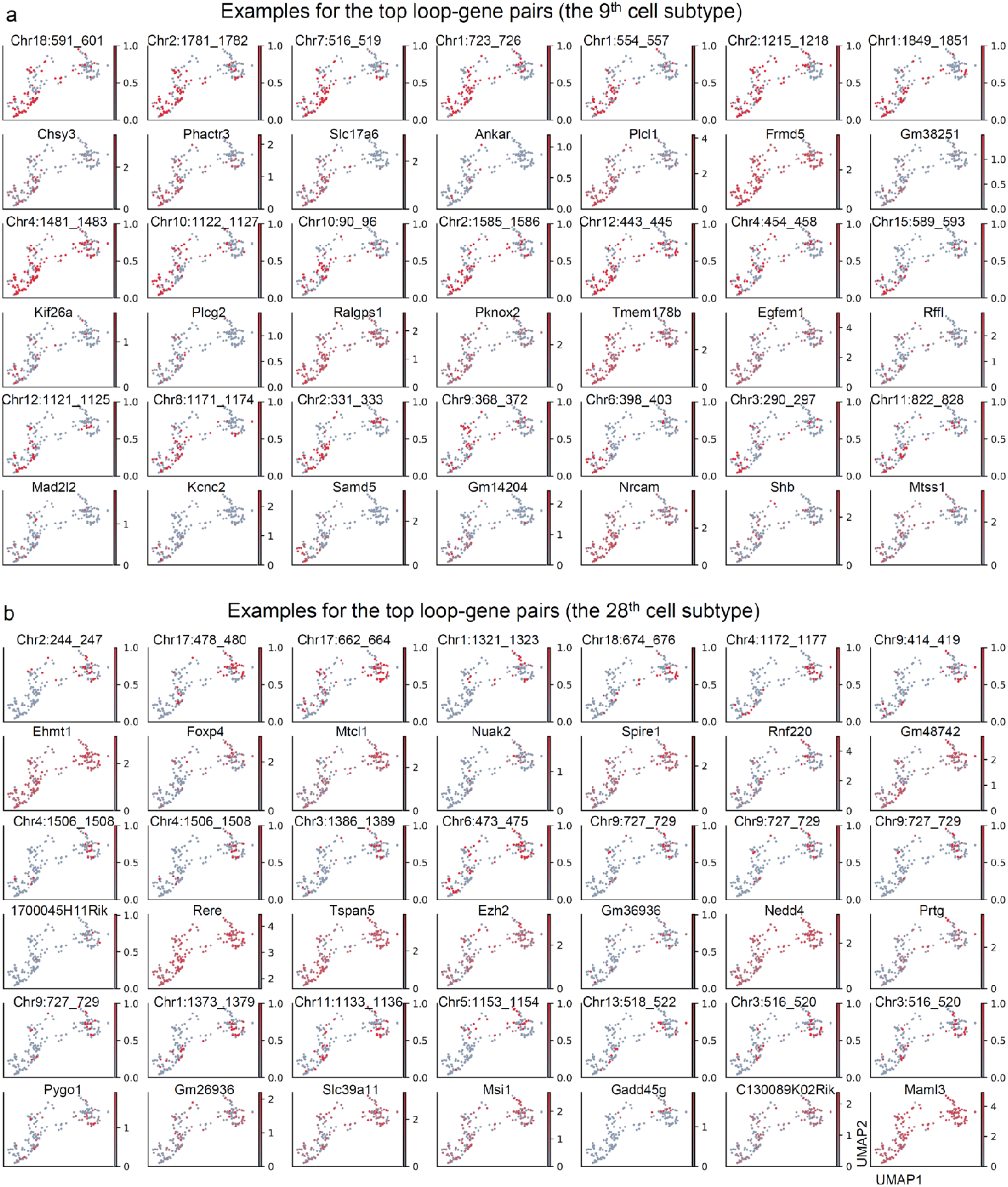
UMAP visualization of the top loop-gene pairs for clusters 9 (a) and 28 (b).

**Extended Data Fig. 10:**
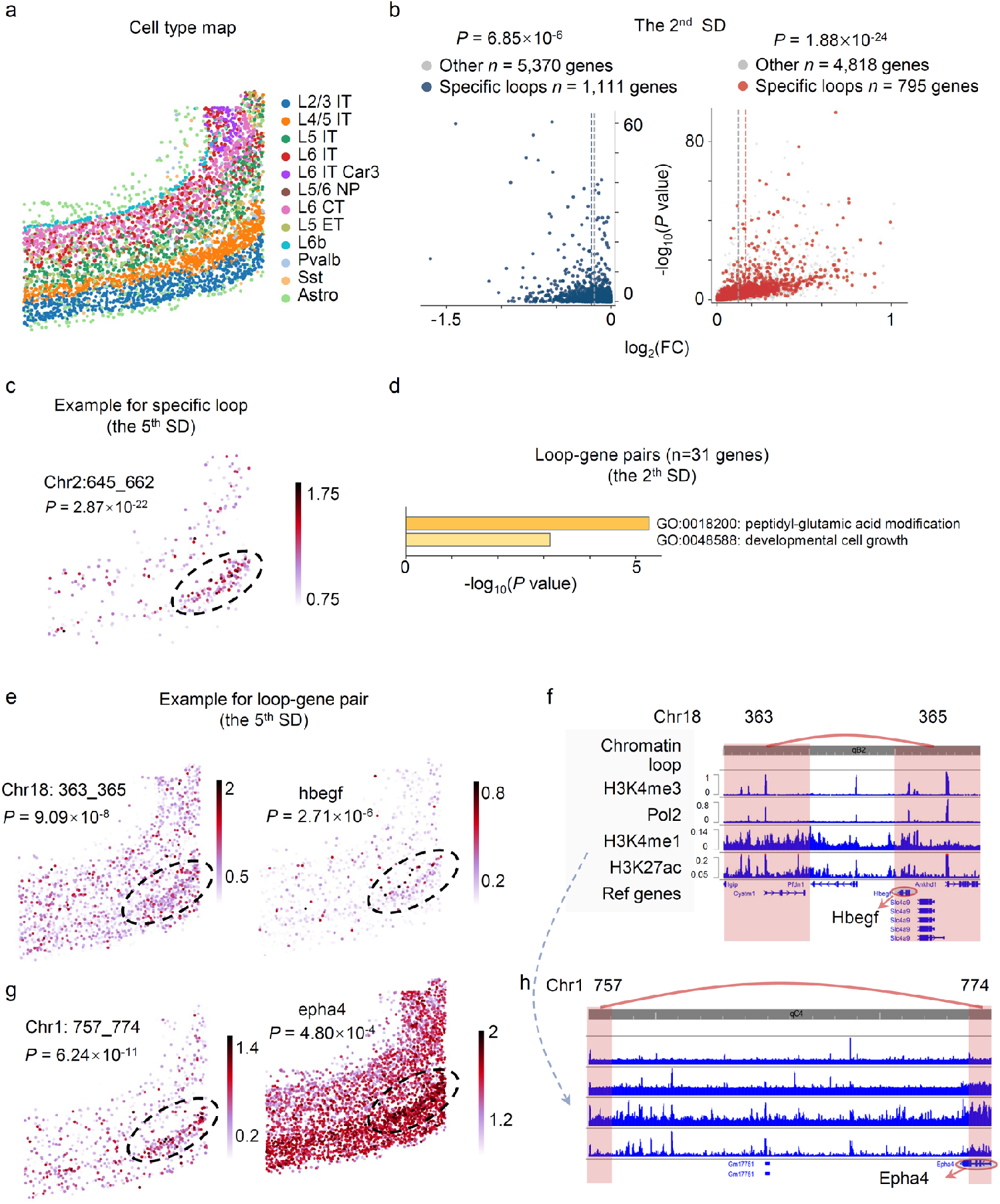
Supplementary explanation for single-cell chromatin loop features depict spatial domains. **a**, in situ plot of cell types. **b**, Left, volcano plot of differential expression. Blue circles denote low-expressed genes within the anchor regions of chromatin loops specific to the 2^th^ SD compared with the rest cells. Grey circles represent other low-expressed genes. Right, Red circles denote high-expressed genes within the anchor regions of chromatin loops specific to the 2^th^ SD. Grey circles represent other high-expressed genes (*P* values from one-sided t-tests). **c**, in situ plot of the example chromatin loop of the 5^th^ SD specific Chr2:645_662. **d**, Gene enrichment analysis of the gene sets associated with loop-gene pairs in the 2^nd^ SD. **e**,**g**, In situ plots of example loop-gene pairs. **f**,**h**, WashU Epigenome Browse shows epigenetic signals around the two chromatin loops, respectively.

