## Supplementary results, methods, figures, and table legends for "CellLoop: Identifying single-cell 3D genome chromatin loops"

**Supplementary Results and Methods**

**Time analysis for identifying chromatin loops of single cells**

The CellLoop algorithm was evaluated on a computing workstation equipped with an Intel(R) Xeon(R) Gold 5220R CPU operating at 2.20 GHz and 256 GB of DDR4 RAM. When running on only one CPU core, CellLoop took an average of 38.3-148s to identify chromatin loops at 10kb resolution for different chromosomes, 15.1-52.9s at 20kb resolution, 6.1-19.5 s at 50kb resolution, and 3.75-11.1s at 100 kb resolution (Supplementary Fig. 1a). We also executed CellLoop in parallel using 10 CPU processes. Notably, CellLoop detected single-cell chromatin loops with an average time of merely 0.99s at 100kb resolution, 1.8s at 50kb resolution, 4.6s at 20kb resolution, and approximately 13s at 10kb resolution. Compared to one CPU process, the parallel implementation in 10 CPU processes boosted the efficiency by 6.5-7.6 times (Supplementary Fig. 1b).

CellLoop detects chromatin loops within a 10 MB range of chromatin via a sliding window approach. Thus, the memory footprint of the one-core computing process does not exceed 2 GB. In future versions of CellLoop, theoretically, all chromosomes of different cells could be computed in parallel. Given sufficient CPU cores and memory space on the computing platform, the operational efficiency of CellLoop will be significantly enhanced.

**The** **hypothesis testing of the overlapping chromatin loops**

The hypergeometric test of the overlapping chromatin loops detected by LFmap and existing methods was conducted using the hypergeom function in the scipy package, relying on the following four numbers: the number of chromatin loops detected by LFmap and existing methods simultaneously; the number of all possible chromatin loops within the maximum distance range; the number of chromatin loops contained in LFmap; the number of chromatin loops detected by existing methods.

We also investigated the hypothesis testing of the chromatin loops in LFmap that overlapped with those detected simultaneously by the SnapHi-C and HiCCUPs methods (Fig. 3e,f). To establish the expected distribution of the chromatin loop number, we carried out sampling without replacement. Precisely, we randomly picked 872 chromatin loops from the 1,192 loops identified by SnapHi-C on chromosome 2, and 217 chromatin loops from the 278 loops detected by HiCCUPs on the same chromosome. Upon repeating this process 100,000 times, we acquired the distribution of the overlapping loop numbers. The significance in terms of *P* value was ascertained by the proportion of the observed overlap count (50) being less than the overlap counts generated from the 100,000 repetitions.

**The selection of the default parameters for the CellLoop algorithm**

The default value of *k* is selected to be in the range of 25 to 50 for linking a single cell to its *k*-nearest neighbors (*k*-NNs). Additionally, we establish another prerequisite for cells to be regarded as neighboring cells, which is that the correlation coefficient associated with the two cells must exceed 0.1. When the parameter *k* is set to its maximum value of 50, in the Dip-C dataset^1^, the median number of neighboring cells is 22. Conversely, when the parameter is set to its minimum value of 25, the median number of actual neighboring cells is 20. Consequently, the magnitude of the parameter *k* has a relatively minor impact on the outcome. A similar phenomenon is also observed in both HiRES^2^ and GAGE-seq^3^ datasets. In practical prototypes, we recommend setting the value of k to 50.

**Normalization of genomic distance for chromatin loops**

The following formula was defined to calculate log2 distance of each chromatin loop:

$$NormD=floor(\frac{{log}_{2}\left( d \right)+s}{s})$$

where $d$ indicates the genomic distance of chromatin loops, $s$ means an exponent step of each bin, whose typical values are 0.25.

**Go-term analysis**

Metascape^4^, a gene annotation or analysis resource (http://metascape.org/gp/index.html#/main/step1), was applied for GO-term enrichment analysis for a given mouse gene set.

**Supplementary Figures**

**Supplementary Fig. 1** | **Time analysis for identifying chromatin loops of single cells. a,** On only one CPU process, running time distribution for identifying single-cell chromatin loops on a single chromosome at different resolutions by CellLoop algorithm. **b,** The average running time for identifying single-cell chromatin loops on a single chromosome at different resolutions comparing a single CPU process with 10 CPU processes.

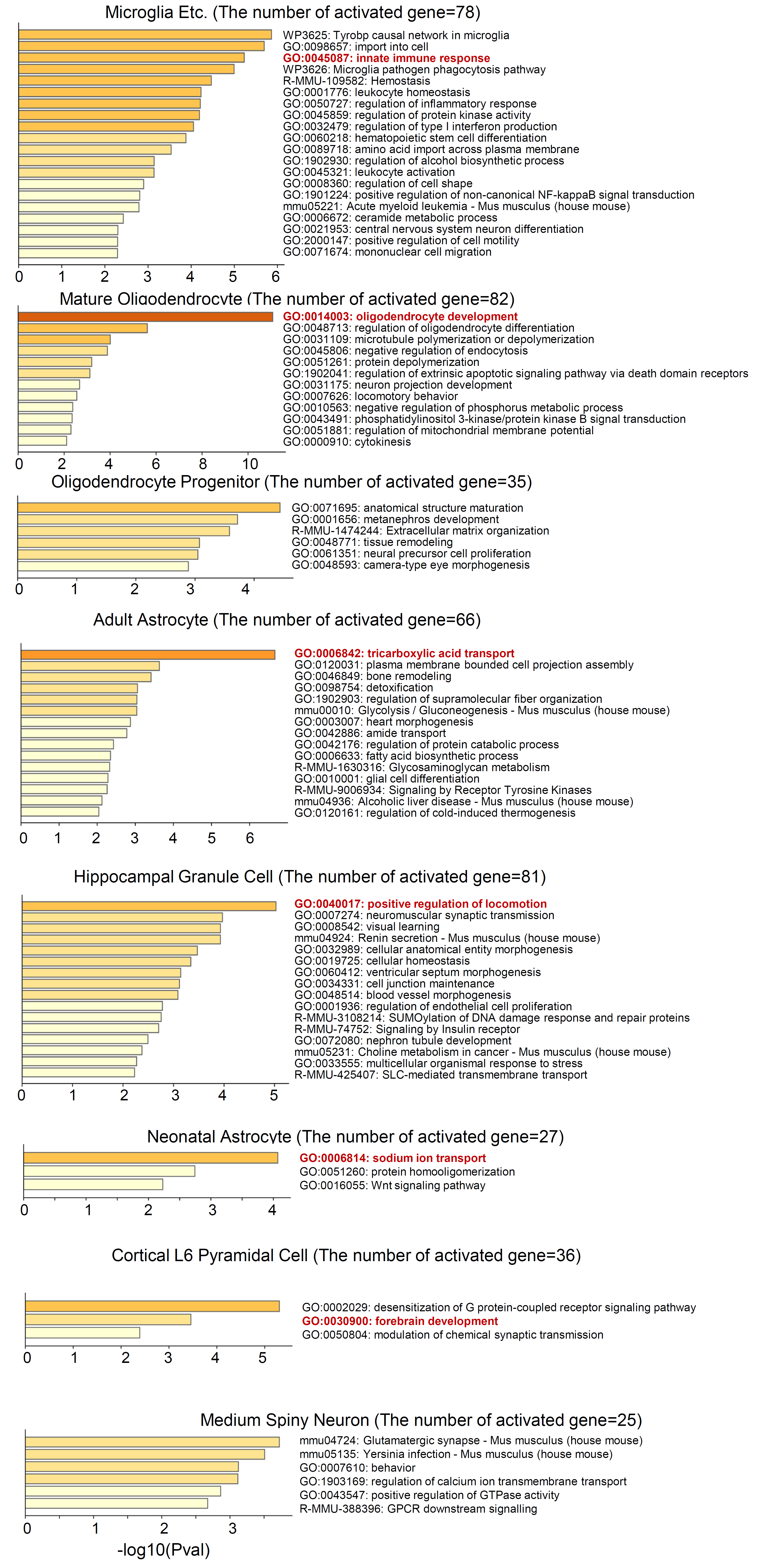

**Supplementary Fig. 2** | **Gene function enrichment analysis for gene sets from Indi-Spe-highgene of different cell types.**

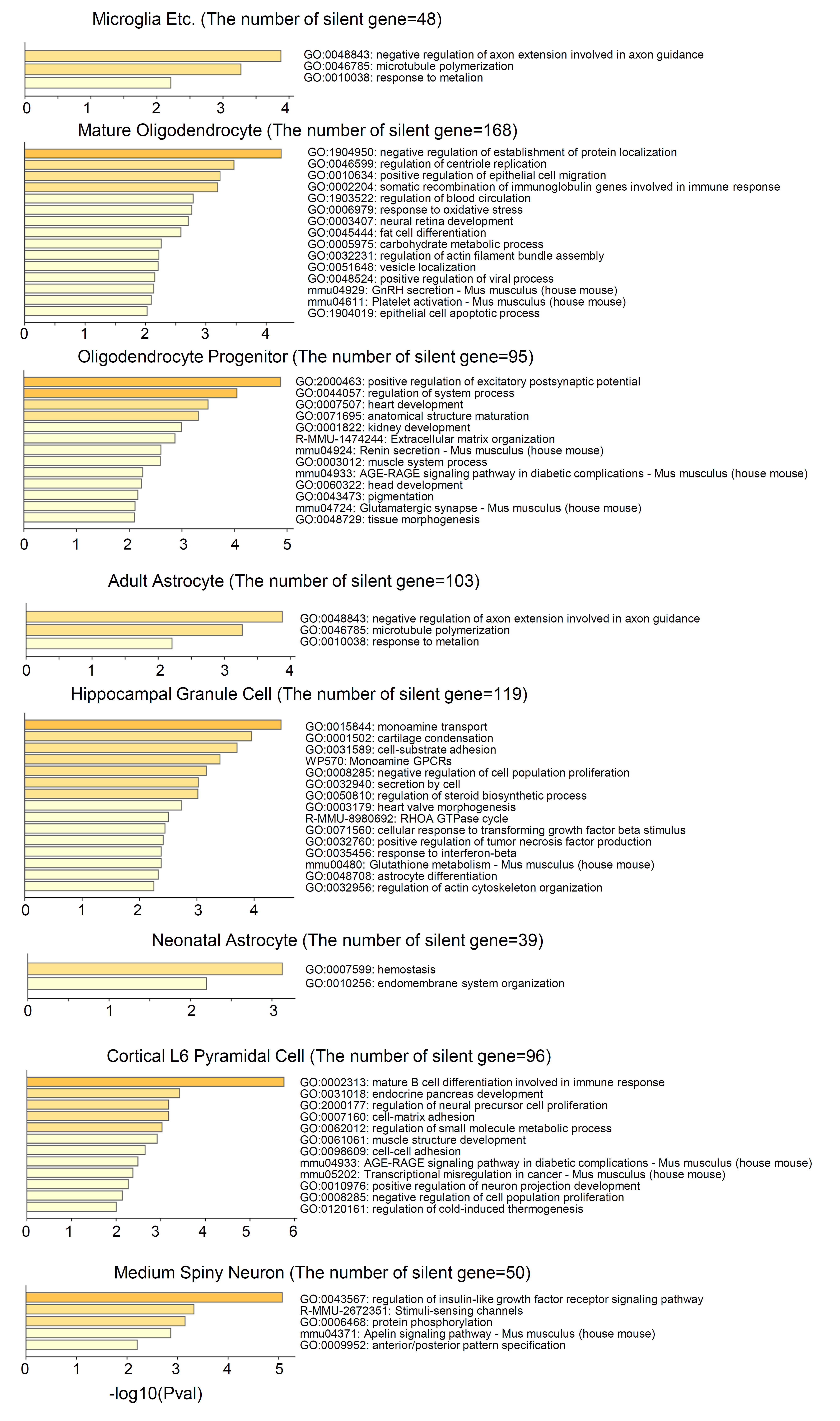

**Supplementary Fig. 3** | **Gene function enrichment analysis for gene sets from Indi-Spe-lowgene of different cell types.**

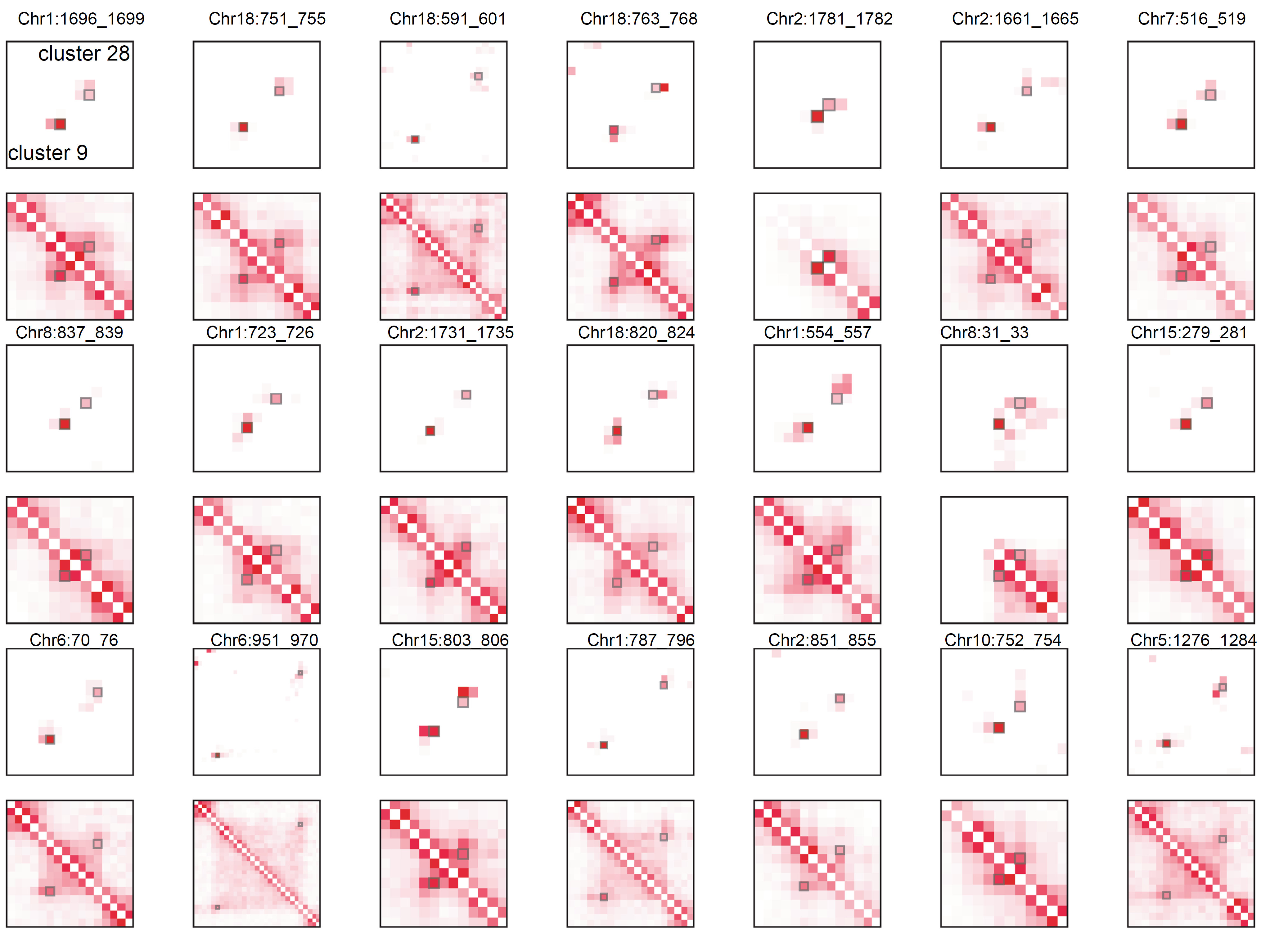

**Supplementary Fig. 4** | **CFmap and LFmap around the top specific chromatin loop of cluster 9 in two clusters.** In all Heatmaps, cluster 9 is at the lower left and cluster 28 at the lower right. Small squares mark specific chromatin loops.

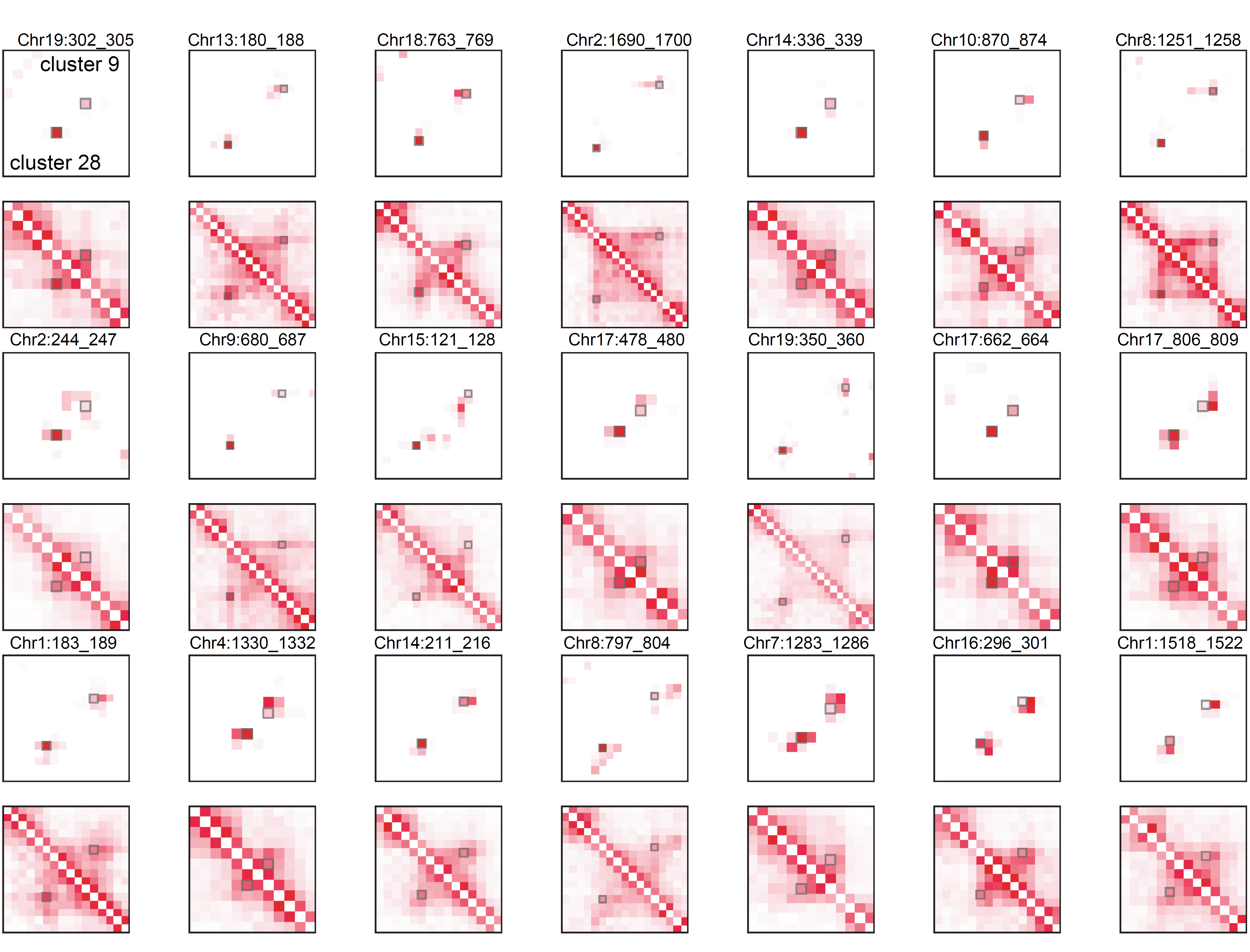

**Supplementary Fig. 5** | **CFmap and LFmap around the top specific chromatin loop of cluster 28 in two clusters.** In all Heatmaps, cluster 28 is at the lower left and cluster 9 at the lower right. Small squares mark specific chromatin loops.

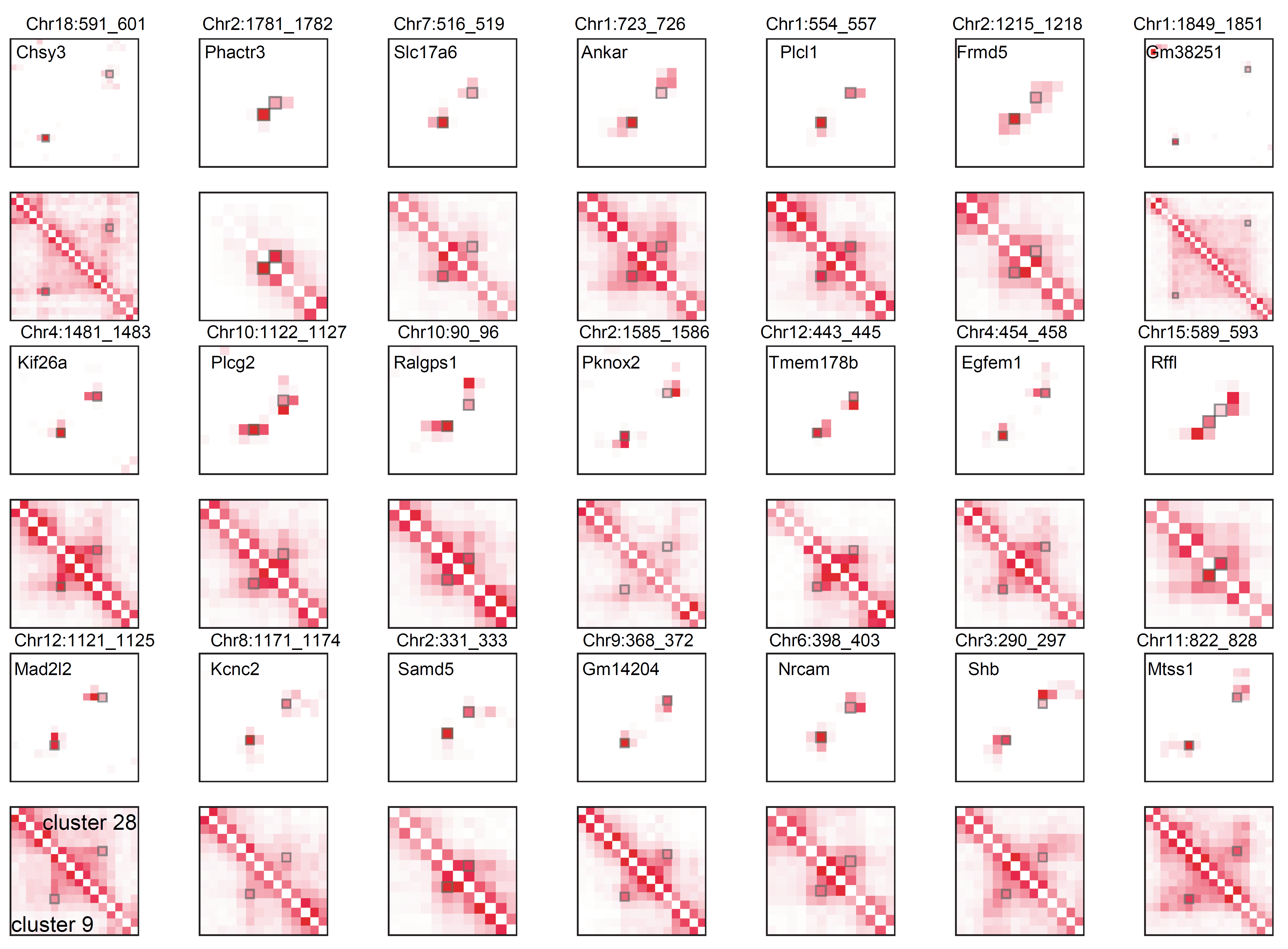

**Supplementary Fig. 6** | **CFmap and LFmap around the top loop-gene pairs of cluster 9 in two clusters.** In all Heatmaps, cluster 9 is at the lower left and cluster 28 at the lower right. Small squares mark specific chromatin loops.

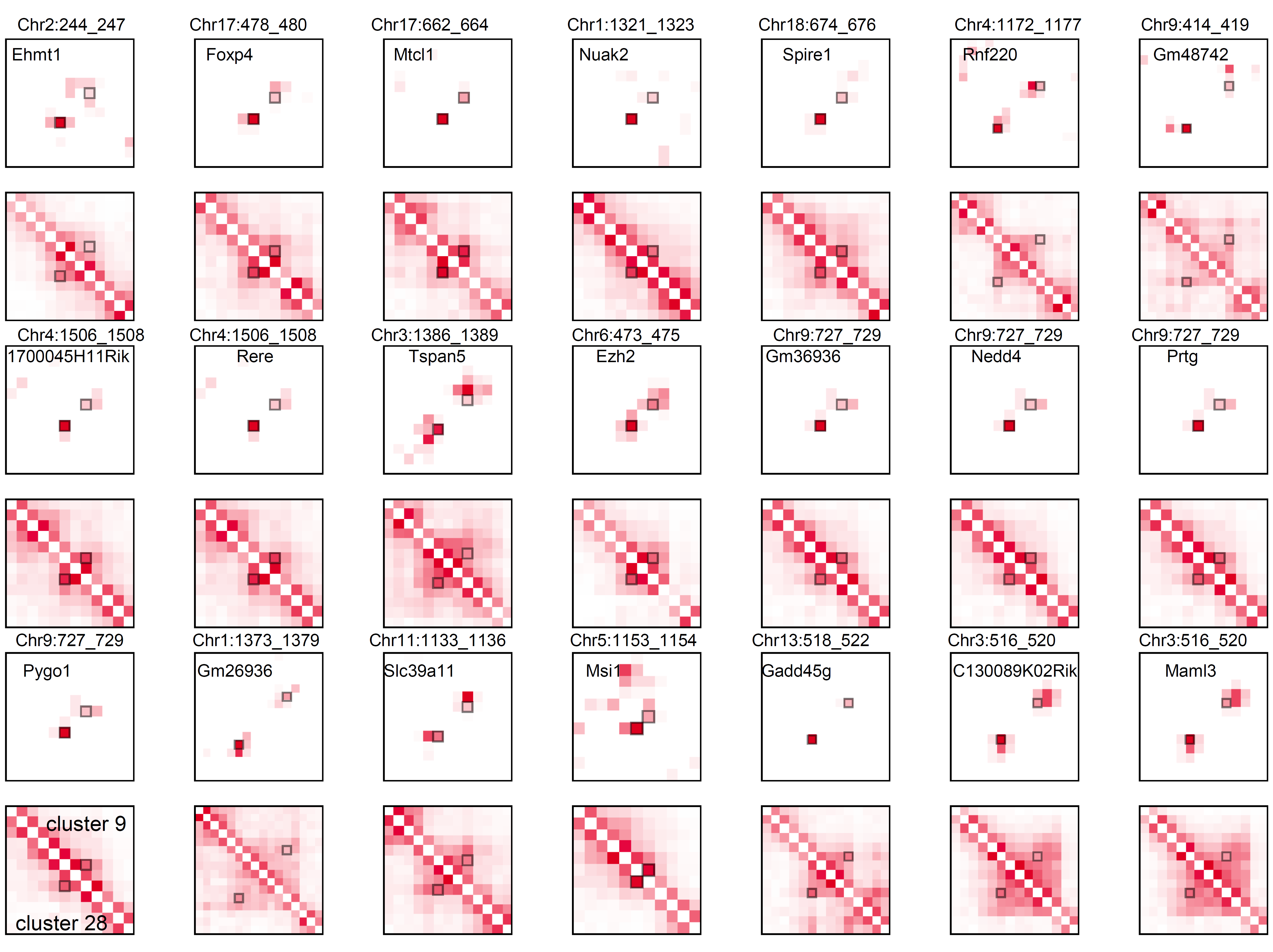

**Supplementary Fig. 7** | **CFmap and LFmap around the top loop-gene pairs of cluster 28 in two clusters.** In all Heatmaps, cluster 28 is at the lower left and cluster 9 at the lower right. Small squares mark specific chromatin loops.

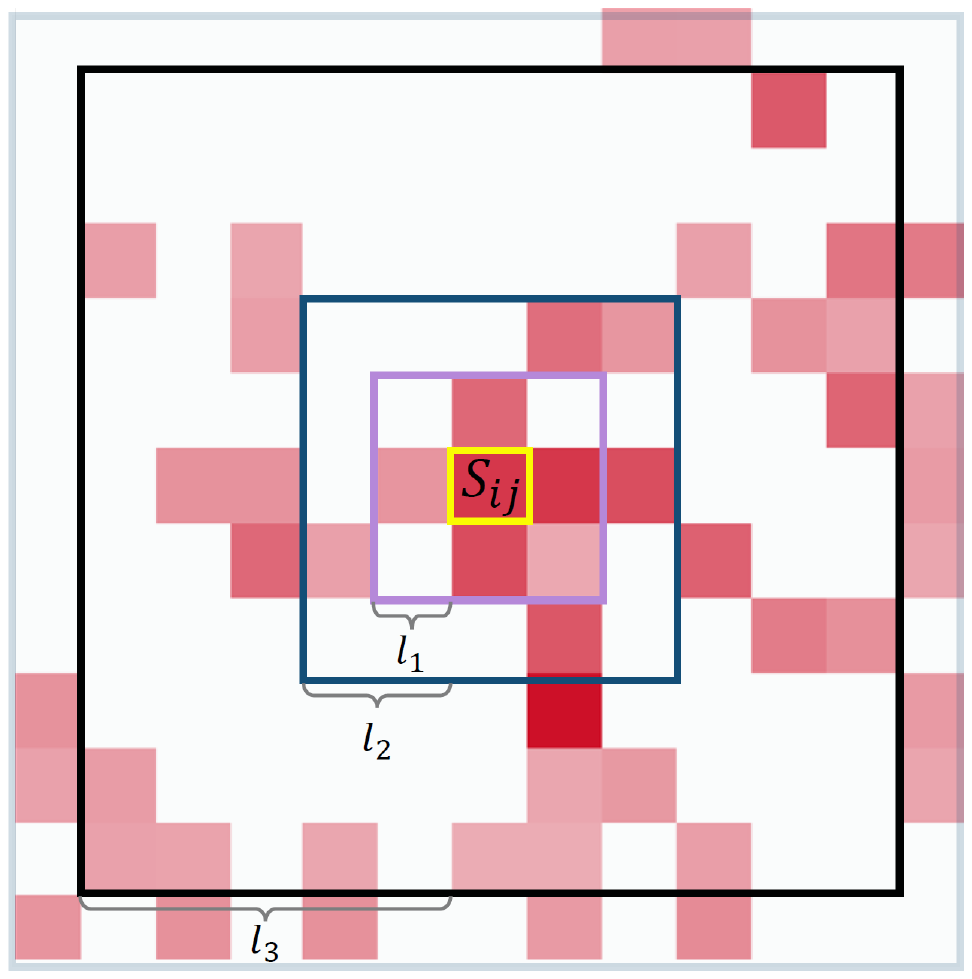

**Supplementary Fig. 8** | **Definition of multiple Parameters in CellLoop.**

**Supplementary Table Legends**

**Supplementary Table 1** | **Gene enrichment analysis of the gene sets associated with loop-gene pairs in the 5^th^ SD.**

**Supplementary Table 2** | **Gene enrichment analysis of the gene sets associated with loop-gene pairs in the 2^nd^ SD.**

**Supplementary Table 3** | **Gene enrichment analysis of the gene sets associated with unassociated loop-gene pairs in the 5^th^ SD.**
